# Metabolomics, Transcriptomics and Functional Glycomics Reveals Bladder Cancer Cells Plasticity and Enhanced Aggressiveness Facing Hypoxia and Glucose Deprivation

**DOI:** 10.1101/2021.02.14.431133

**Authors:** Andreia Peixoto, Rui Freitas, Dylan Ferreira, Marta Relvas-Santos, Paula Paulo, Marta Cardoso, Janine Soares, Cristiana Gaiteiro, Carlos Palmeira, Filipe Teixeira, Rita Ferreira, Maria José Oliveira, André M. N. Silva, Lúcio Lara Santos, José Alexandre Ferreira

**Author notes:** **Corresponding author:** José Alexandre Ferreira, Experimental Pathology and Therapeutics Group, Research Centre, Portuguese Oncology, Institute of Porto, R. Dr. António Bernardino de Almeida 62, 4200-072 Porto, Portugal; (ext. 5111).

## Abstract

Bladder cancer constitutes one of the deadliest genitourinary diseases, especially when diagnosed at late stages. These tumours harbour microenvironmental niches characterized by low levels of oxygen (hypoxia) and limited glucose supply due to poor vascularization. However, the synergic contribution of these features to disease development is poorly understood. Here, we demonstrated that cells with distinct histopathological and molecular backgrounds responded similarly to such stimuli. Cancer cells arrested proliferation, significantly increased invasive capacity *in vitro* and enhanced tolerance to cisplatin-based chemotherapy. Reoxygenation and access to glucose restored basal proliferation and invasion levels without triggering stress-induced apoptosis, denoting significant cellular plasticity in adapting to microenvironmental cues. Whole transcriptomics showed major molecular reprogramming, supporting main functional alterations. Metabolomics evidenced fatty acids *β*-oxidation as main bioenergetic pathway rather than anaerobic glycolysis generally adopted by hypoxic cells. Joint pathway analysis also suggested relevant alterations in mucin-type *O*-glycan biosynthesis. Glycomics confirmed a major antagonization of *O*-glycosylation pathways, leading to simple cell glycophenotypes characterized by the accumulation of immature short-chain *O*-glycans such as Tn and STn antigens at the cell surface. Glycoengineered models reflecting simple cell glycophenotypes were developed and functional studies *in vitro* and *in vivo* showed that Tn and STn overexpression decreased proliferation and promoted chemoresistance, reinforcing their close link with tumour aggressiveness. Collectively, we have demonstrated that hypoxia and glucose deprivation trigger more aggressive cell behaviours, in what appears to be an escape mechanism from microenvironmental stress. We propose that, altered glycosylation may be used to target these subpopulations, paving the way for precision oncology.

## 1. Introduction

Bladder cancer (BC) remains one of the deadliest malignancies of the genitourinary tract due to high intra and inter-tumoral molecular heterogeneity. This has delayed a more comprehensive understanding on tumour spatiotemporal evaluation and affected the efficiency of precise clinical interventions. While genetic alterations are considered primary causes of cancer development, downstream phenotypic changes induced by the tumour microenvironment are amongst the driving forces of progression and dissemination. The generation of hypoxic niches characterized by decreased oxygen availability (≤2% O_2_) is a microenvironment hallmark of solid tumours^1^. Not surprisingly, the presence of hypoxic regions is a pivotal independent poor prognosis factor in several cancers, including urothelial carcinomas^2,3^.

Uncontrolled tumour cell proliferation supported by avid glucose consumption and glycolysis in the presence of oxygen and fully functioning mitochondria (the Warburg effect) are common features of solid tumours^4^. Rapid proliferation is also frequently accompanied by flawed neoangiogenesis, resulting in suboptimal oxygen and nutrients supply to cancer cells in the periphery of blood vessels. Moreover, poor vascularization and competition for nutrients requires constant metabolic remodelling and exploitation of alternative survival strategies by hypoxic and nutrient deprived cells, such as the induction of cellular quiescence^5^. While many tumour cells faced with suboptimal growth conditions undergo programmed cell death and necrosis, some subpopulations show tremendous molecular plasticity adapting to hypoxic and nutrient deprived microenvironments^6^. Namely, more plastic tumour cells frequently undergo a massive metabolic reprograming towards anaerobic glycolysis and mitochondrial autophagy accompanied by an increase in lactate biosynthesis and tumour microenvironment acidification, with major implications in cancer progression^7,8^. The selective pressure caused by stress factors further contributes to the maintenance of cancer stem cells and promotes the acquisition of epithelial-to-mesenchymal transition (EMT) traits that decisively dictate tumour fate^9^. Moreover, slow dividing bladder cancer cells in hypoxic regions can escape many cytotoxic drugs targeting rapidly dividing cells and are also sufficiently shielded from many other therapeutic agents when compared to the tumour bulk^10^. However, the exact molecular mechanisms by which oxygen gradients and glucose deprivation induce more aggressive and metastatic phenotypes remain poorly explored in bladder cancer. Furthermore, there is little knowledge on molecular alterations occurring at the cell surface which dictate poor prognosis and may be easily targeted in theragnostic interventions.

The cell surface is densely covered by an extended layer of complex glycans and glycoconjugates of different natures. Glycans are not direct gene products, but rather the concerted result of a wide variety of glycosyltransferases, glycosidases, and sugars nucleotide transporters across the secretory pathways^11^. Moreover, glycosylation enables rapid accommodation of microenvironmental stimuli in response to alterations in glycogenes expression and metabolic imbalances^12^. These alterations directly influence protein trafficking, stability, and folding and decisively shape the cancer cells proteome with impact on all cancer hallmarks^1^. These include activation of oncogenic signalling transduction, induction of immune tolerance, migration, cell-cell, and cell-matrix adhesion^1^. In early reports, we have suggested that hypoxia may significantly antagonize protein glycosylation of serine and threonine residues (*O*-glycosylation) and concomitantly induce EMT, responsible by increased cell invasion^12^. The cancer-associated short-chain *O*-glycan Sialyl-Tn (STn) was the most prominent glycan arising from these alterations, being associated with immune tolerance and worst prognosis in bladder cancer^12–14^. However, a direct link between the glycan and cell invasion in this context needs further demonstration. The influence of glucose was also not estimated, requiring a more in-depth functional characterization of the glycome facing different microenvironments.

In this study, we exploit an integrated multi-omics approach combining metabolomics and whole transcriptomics to gain insights on the molecular plasticity of bladder cancer cells facing hypoxia and low glucose. We further devote to understanding the effect of the microenvironment across the glycosylation axis. Functional glycomics supported by a library of well characterized glycoengineered cell models was used to determine how altered glycosylation contributes to bladder cancer progression. Important insights were generated to identify and address more aggressive bladder cancer subpopulations in precision oncology settings.

## 2. Material and Methods

### 2.1. Cell culture conditions

Human BC cell lines RT4, 5637, T24, and HT1197 were purchased from American Type Culture Collection (ATCC). RT4, 5637, and T24 cells were maintained with complete RPMI 1640 GlutaMAX™ medium (Gibco) and HT1197 with DMEM GlutaMAX™ medium (Gibco). Cells were kept at 37 °C in a 5% CO_2_ humidified atmosphere (normoxia). Cells were also grown under hypoxia and nutrient deprivation at 37°C in a 5% CO_2_, 99.9% N_2_ and 0.1% O_2_ atmosphere using a BINDER C-150 incubator (BINDER GmbH) and complete RPMI 1640 and DMEM media without glucose (Gibco). In re-oxygenation experiments, cells under oxygen and glucose deprivation were restored to standard culture conditions 24h prior to analysis.

### 2.2. CRISPR-Cas9 glycoengineered cell models

A recombinant *Streptococcus pyogenes* Cas9 (GeneArt^TM^ Platinum Cas9 Nuclease, Thermo Fisher Scientific) together with a single-guided RNA (GTAAAGCAGGGCTACATGAG, sgRNA) were used to generate site-specific double-strand breaks (DSBs) in the *C1GALT1* gene in T24 cells *in vitro*. In parallel, two sgRNAs were used for *GCNT1* gene knock-out (KO) (gRNA1: TAGTCGTCAGGTGTCCACCG, gRNA2: AAGCGGTATGAGGTCGTTAA). Lipofectamine™ CRISPRMAX™ Transfection Reagent (Thermo Fisher Scientific) was used according to the manufacturer’s instructions. Complexes were made in serum-free media (Opti-MEM™ I Reduced Serum Medium) and added directly to cells in culture medium and incubated for 24 h. Single clones were obtained by serial dilution in 96 well plates and KO clones were identified by Indel Detection by Amplicon Analysis (IDAA) using ABI PRISM™ 3010 Genetic Analyzer (Thermo Fisher Scientific) and Sanger sequencing. Three clones were selected for each gene with distinct out of frame indel formation. Single clones with silent mutations provided phenotypic control cell lines. IDAA results were analysed using Peak Scanner Software V1.0 (Thermo Fisher Scientific). Human *ST6GALNAC1* (hST6GALNAC1 [NM_018414.5]) knock-in (KI) was performed in C1GALT1 KO cells by conventional mammalian gene expression vector transfection, using jetPRIME® transfection reagent (PolyPlus Transfection) according to product instructions. Positively transfected cells were selected based on puromycin (2µg/mL, EMD Millipore) resistance. In parallel, a mock system containing a 300 bp stuffer ORF was developed.

### 2.3. Cell apoptosis assay

Apoptosis was determined using the Cell Apoptosis Kit with FITC annexin V and PI for flow cytometry (Thermo Fisher Scientific) according to the manufacturer’s instructions. Briefly, cells cultured under normoxia and hypoxia plus glucose deprivation were detached using Accutase enzyme cell detachment medium (Thermo Fisher Scientific) and stained with recombinant annexin V conjugated to fluorescein (FITC annexin V) as well as with red-fluorescent propidium iodide (PI) nucleic acid binding dye. Data analysis was performed through CXP Software in a FC500 Beckman Coulter flow cytometer.

### 2.4. Flow cytometry for short-chain *O*-glycans

Cells were detached using Versene solution (Thermo Fisher Scientific), fixed with 1% paraformaldehyde (PFA; Sigma-Aldrich) and stained with mouse anti-TAG72 (B72.3+CC49) (Abcam) using a 2 μg/10^6^ cells dilution in PBS 2% FBS for 1 h at room temperature. Polyclonal rabbit anti-mouse immunoglobulins/FITC (DAKO) were used as secondary antibodies for STn detection at a 1:100 dilution in PBS 2% FBS for 15 minutes at room temperature. Mouse IgG1 [MOPC-21] isotype control (Abcam) was included as negative control. In parallel, 10^6^ cells were digested with 70 mU Neuraminidase from *Clostridium perfringens* (Sigma-Aldrich) in sodium acetate buffer at 37 °C overnight, under mild agitation, prior to STn staining, and used as negative controls. In addition, cancer cells were screened for Tn and T antigens as well as for *N*-acetylglucosamine residues using fluorescein-labelled lectins (Vector Laboratories) as *Vicia Villosa* (VVA, 0.01 mg/mL), Peanut Agglutinin (PNA, 0.01 mg/mL) and *Griffonia Simplicifolia* Lectin II (GSL II, 0.02 mg/mL), respectively. VVA and PNA lectins were incubated for 1 h in PBS 2% FBS under mild agitation at room temperature, while the lectin GSL II was incubated in 10 mM HEPES, 0.15 M NaCl, 0.1 mM CaCl_2_, pH 7.5 buffer. Sialylated Tn and T antigens expression was determined after neuraminidase treatment under the above-mentioned conditions. GSL II lectin detection was performed following PNGaseF (250 mU/10^6^ cells, in PBS 1x; Sigma-Aldrich) enzymatic digestion to exclude *N*-glycan-associated *N*-acetylglucosamine residues contribution. Data analysis was performed through CXP Software in a FC500 Beckman Coulter flow cytometer.

### 2.5. Immunofluorescence for short-chain *O*-glycans detection

To evaluate short *O*-glycans expressions, T24 glycoengineered cell models were cultured at low density and fixed with 4% paraformaldehyde (PFA, Sigma-Aldrich), following immunofluorescent staining similar to the flow cytometry protocol. Sialylated glycoforms were evaluated in parallel with samples digested with 50 mU/mL α-neuraminidase from *Clostridium perfringens* for 4 h at 37 °C. After antigen staining, cells were marked with 2,3×10^−3^ μg/μL 4’,6-Diamidino-2-phenylindole dihydrochloride (DAPI, Thermo Fisher Scientific) for 10 minutes at room temperature in the dark. All images were acquired on a Leica DMI6000 FFW microscope using Las X software (Leica).

### 2.6. Cell proliferation assay

Cell proliferation was evaluated using a colorimetric Cell Proliferation ELISA (Roche), based on the measurement of the incorporation of bromodeoxyuridine (BrdU) into newly synthesized DNA of proliferative cells. Procedure steps were followed according to the manufacturer instructions and results were monitored at 450 nm using a microplate reader (iMARK™, Bio-Rad).

### 2.7. Cell cycle analysis

Cells were harvested by trypsinization, following fixation in cold 70% ethanol for 30 minutes at 4°C. After washing, cells were resuspended in 1mL of ready to use DNA labelling solution (Cytognos)/10^6^ cells and incubated in horizontal position for 10 minutes at room temperature in obscurity. Data analysis was performed through CXP Software in a FC500 Beckman Coulter flow cytometer.

### 2.8. Invasion Assays

Invasion assays were performed under normoxia and hypoxia plus glycose deprivation using Corning® BioCoat™ Matrigel® Invasion Chambers as described in Peixoto, A. *et al*.^12^. Invasion assays were normalized to cells proliferation and cells were seeded in quintuplicates for each experiment. Gelatine zymography was performed using conditioned media from invasion assays to determine proteolytic activity of matrix metalloproteinases (MMP) 2 and 9 under the experimental conditions as described in Peixoto, A. *et al*.^12^.

### 2.9. L-lactate Assay

An L–Lactate colorimetric Assay Kit (Abcam) was used to detect L(+)-Lactate in deproteinized cultured cells lysates and conditioned medium. Procedure steps were followed according to the manufacturer’s instructions and results were monitored at 450 nm using a microplate reader (iMARK™, Bio-Rad). The results were normalized to cell proliferation.

### 2.10. Western Blot

Whole protein extracts were collected from bladder cancer cells using a 25 mM Tris-HCl, pH 7.2, 150 mM NaCl, 5 mM MgCl_2_, 1% NP-40, and 5% glycerol lysis buffer, supplemented with Halt™ Protease and Phosphatase Inhibitor Cocktail (Thermo Fisher Scientific). Twenty micrograms of isolated proteins were run on 4–20% precast SDS-PAGE gels (BioRad), transferred into nitrocellulose membranes, and screened using a mouse monoclonal antibody to AMPK alpha 1 + AMPK alpha 2 (Abcam), 1:1000 for 1 h at room temperature. Peroxidase AffiniPure Goat Anti-Mouse IgG (Jackson ImmunoResearch) was used as secondary antibody at 1:70 000 for 30 minutes at room temperature. A Rabbit polyclonal antibody to AMPK alpha 1 (phospho T183) + AMPK alpha 2 (phospho T172) (Abcam) was also used at 1:1000 dilution for 1h at room temperature as well as the respective peroxidase conjugated goat anti-Rabbit IgG secondary antibody (ThermoFisherScientific) at 1:60 000 for 30 minutes at room temperature. A rabbit monoclonal antibody to beta 2 Microglobulin (Abcam) was used as loading control. Total protein stain was also assessed using Ponceau S (BioRad) anionic dye.

### 2.11. HIF-1α expression

Hypoxia-inducible factor 1-alpha (HIF-1α) was evaluated using a HIF-1 Alpha ELISA Kit (Invitrogen™). Procedure steps were followed according to the manufacturer’s instructions and results were monitored at 450 nm using a microplate reader (iMARK™, Bio-Rad). The results were normalized to cell proliferation.

### 2.12. Metabolomics

Cells were dispersed in 80% methanol (Merck), sonicated for 30 minutes at 4°C and kept at −20°C for 1 h. Samples were then centrifuged, and the supernatant was analysed by UHPLC-ESI-MS/MS in positive and negative mode. Metabolite analysis was performed using an Ultimate 3000LC combined with Q Exactive mass spectrometer (Thermo Fisher Scientific). Eluent A was 0.1% formic acid in water and eluent B was acetonitrile and metabolite separation occurred using the following gradient elution (0-1.5 min, 95-70% A; 1.5-9.5 min, 70-5% A; 9.5-14.5 min, 5% A; 14.5-14.6 min, 5-95% A; 14.6-18.0 min, 95% A). The flow rate of the mobile phase was 0.3 mL/min. The column (Acquity UPLC HSS T3; 100 Å, 1.8 μm, 2.1 mm × 150 mm) temperature was maintained at 40°C, and the sample manager temperature was set at 4°C. Metabolites were identified by retention time and corresponding MS/MS spectra. For metabolomics data pre-processing and analysis, raw data matrices were blank subtracted (a mean blank value was calculated per metabolite) and normalized to the number of cells for each condition. The resulting matrices were then imported to Metaboanalyst 4.0 and log-transformed to reduce heteroscedasticity and pareto-scaled to adjust for differences in fold-changes between metabolites.

### 2.13. Metabolomics data analysis

Multivariate and univariate analysis were performed to identify metabolites that discriminate normoxia from hypoxia with low glucose. Unsupervised principal components analysis (PCA) was applied to unravel data structure, following a supervised method, namely partial-least-squares discriminant analysis (PLS-DA) to identify which metabolites are useful to predict group membership. Metabolites with discriminative power were ranked based on Variable Importance in Projection (VIP) values >1 and PLS-DA models were validated based on the “prediction accuracy during training” test statistic with 1000 permutations (*p*<0.05 for significance). Heat maps with hierarchical clustering of metabolites were constructed based on the following metrics: i) distance measure: Pearson correlation (similarity of expression profiles), ii) clustering algorithm: complete linkage (forms compact clusters), iii) feature autoscale. Hierarchical clustering of samples was carried out based on the following metrics: i) distance measure: Euclidean distance (sensitive to magnitude differences), ii) clustering algorithm: Ward (minimizes within-cluster variance). Differences in metabolites between groups were further evaluated using one-way analysis of variance (ANOVA) with a False Discovery Rate (FDR) cut-off set at 0.05 for significance. Tukey’s post hoc was applied to check which groups differed. Significant metabolites unravelled by ANOVA were then used for pathway analysis to identify the most relevant pathways that are involved in the adaptation of cells from normoxia to hypoxia with low glucose. Pathway analysis was carried out based on two features: i) functional enrichment which was assessed using hypergeometric test for over-representation analysis (*p*<0.05 for significance) and ii) pathway topology analysis, which was implemented using the relative betweenness centrality. Pathway impact was considered relevant if > 0.1. The joint pathway analysis was carried out using transcriptomic and metabolomic data, based on a gene and metabolite list with associated fold-changes. The human pathway library was chosen using the pathway database “all pathways (integrated)”. The enrichment analysis was based on the hypergeometric test statistic while degree centrality was used as topology measure. The integration method was based on a combination of queries.

### 2.14. Citrate synthase assay

Citrate synthase (CS) activity was measured in whole cell protein lysates using the method proposed by Coore et al. (1971)^15^. In brief, the CoASH released from the reaction of acetyl-CoA (Sigma-Aldrich) with oxaloacetate (OAA, Sigma-Aldrich) was determined by its reaction with 5,5’-dithiobis-(2-nitrobenzoic acid) (DTNB, Sigma-Aldrich) at 412 nm (ε=13.6 mM^−1^cm^−1^) in a microplate reader (iMARK™, Bio-Rad). The results were normalized to cell proliferation.

### 2.15. ATP detection assay

A fluorometric ATP assay kit (Abcam) was used according to the manufacturer’s instructions to determine ATP levels in deproteinized whole cell lysates. The ATP assay protocol relied on the phosphorylation of glycerol to generate a fluorometric product (Ex/Em = 535/587 nm), which was quantified using a Synergy™ Mx microplate reader. The results were normalized to cell proliferation.

### 2.16. Transcriptomics

Total RNA was extracted from cell pellets using the RNeasy Plus Mini kit (Qiagen). RNA samples were quantified using Qubit 2.0 Fluorometer (Life Technologies) and RNA integrity was checked with Agilent TapeStation (Agilent Technologies). RNA sequencing library preparations were performed using NEBNext Ultra RNA Library Prep Kit for Illumina following manufacturer’s recommendations (New England Biolabs). Briefly, mRNAs were first enriched with Oligod(T) and fragmented for 15 minutes at 94°C. First strand and second strand cDNA were subsequently synthesized. cDNA fragments were end-repaired and adenylated at 3’ends, and universal adapters were ligated to cDNA fragments, followed by index addition and library enrichment with limited cycle PCR. The sequencing libraries were validated on the Agilent TapeStation (Agilent Technologies) and quantified using Qubit 2.0 Fluorometer (Invitrogen) as well as by quantitative PCR (KAPA Biosystems). The sequencing libraries were clustered on one lane of a flow cell. After clustering, the flow cell was loaded on the Illumina HiSeq 4000 instrument according to manufacturer’s instructions. The samples were sequenced using a 2×150 Paired End (PE) configuration. Image analysis and base calling were conducted by the HiSeq Control Software (HCS). Raw sequence data (.bcl files) generated from Illumina HiSeq was converted into fastq files and de-multiplexed using Illumina's bcl2fastq 2.17 software. One mismatch was allowed for index sequence identification. After investigating the quality of the raw data, sequence reads were trimmed to remove possible adapter sequences and nucleotides with poor quality using Trimmomatic v.0.36. The trimmed reads were mapped to the human reference genome available on ENSEMBL using the STAR aligner v.2.5.2b. Unique gene hit counts were calculated by using feature Counts from the Subread package v.1.5.2. Only unique reads that fell within exon regions were strand-specifically counted. After extraction of gene hit counts, a SNP/INDEL analysis was performed using mpileup within the Samtools v.1.3.1 program followed by VarScan v.2.3.9. The parameters for variant calling were minimum frequency of 25%, *p*-value less than 0.05, minimum coverage of 10, minimum read count of 7. A gene fusion analysis was performed using STAR Fusion v.1.1.0. For novel transcript discovery, transcripts expressed in each sample were extracted from the mapped bam files using Stringtie. The resulting gtf file was compared to the reference annotation file and novel transcripts were identified.

### 2.17. *O*-glycomics

Bladder cancer cellular models *O*-glycome was characterized through the Cellular *O*-glycome Reporter/Amplification method^16,17^. Briefly, benzyl 2-acetamido-2-deoxy-α-D-galactopyranoside (Sigma-Aldrich) was peracetylated and administered to semi-confluent bladder cancer cells, as previously described in Fernandes, E. *et al*.^17^. Secreted benzyl-*O*-glycans were recovered from cell culture media by filtration and solid-phase extraction with a C18 reverse phase sorbent. Finally, Bn-*O*-glycans were permethylated and analysed by reverse phase nanoLC-ESI-MS/MS, as previously described by us^17^. *O*-glycans structures represented in spectra are proposed structures, considering previous knowledge on bladder cancer *O*-glycosylation, chromatography retention times, *m/z* identification, and corresponding product ion spectra.

### 2.18. Anchorage-independent growth

AIG was measured using the soft agar colony formation assay. A 0.5% low melting point agarose (Lonza) solution in complete medium was used as bottom layer in 6-well flat bottom plates (Falcon®). A top layer of 0.3% agarose containing 1×10^4^ cells was then plated and covered with culture medium. Cells were maintained in standard growth conditions for one month. Colonies were then fixed with 10% neutral buffered formalin solution (Sigma-Aldrich) and stained with 0.01% (w/v) crystal violet (Sigma-Aldrich) for 60 minutes. Colonies were photographed using a stereomicroscope (Olympus, SZX16 coupled with a DP71 camera) and automatically counted using the open-source software ImageJ (Fiji package). Only colonies containing more than 50 cells were considered.

### 2.19. Cisplatin resistance assays

Bladder cancer cells were plated into 96 well plates, following a 24 h exposure to crescent concentrations of cisplatin. Positive and negative controls of cell death were set, consisting of cell incubation with complete medium with and without 1% Triton-X (Sigma-Aldrich), respectively. After cisplatin incubation, conditioned media was replaced by a 1.2 mM 3-(4,5-dimethylthiazol-2-yl)-2,5-diphenyltetrazolium bromide (MTT, Thermo Fisher Scientific) solution, following a 4 h incubation at 37°C in a humidified chamber. Finally, formazan crystals were solubilized with dimethyl sulfoxide (DMSO, Sigma-Aldrich) and plates were incubated at 37°C for 10 minutes, following well absorbance measurement at 540 nm in a microplate reader (iMARK™, Bio-Rad). The percentage of cell viability was calculated as follows:

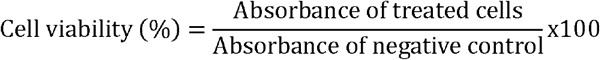

### 2.20. Chicken embryo chorioallantoic membrane (CAM) assay

The chicken embryo CAM assay was used to assess the *in vivo* establishment of tumour aggregates derived from glycoengineered cell models. On embryonic development day (EDD) 3, a squared window was opened in the eggshell of fertilized chicken (*gallus gallus*) eggs, and 2–2.5 mL of albumen was removed to allow detachment of the developing CAM. The window was then sealed with adhesive tape and the eggs were incubated horizontally at 37.8°C in a humidified atmosphere. On EDD9, 1×10^6^ cells derived from each developed cell model were re-suspended in 10 μl of Corning® Matrigel® Matrix and placed in a 3 mm silicone ring attached to the growing CAM. Control cells and respective glycoengineered models were inoculated in the same egg, at least 10 viable embryos were used per experimental pair. The eggs were re-sealed and returned to the incubator for one week. On EDD16, the CAM was excised from the embryos, photographed *ex-ovo* under a stereoscope at 20x magnification (Olympus, SZX16 coupled with a DP71 camera), and images were analysed to determine tumour size. CAM attached tumours were then formalin fixed and paraffin embedded, following haematoxylin and eosin staining of selected samples.

### 2.21. Transmission Electron Microscopy

For electron microscopy, T24 and 5637 cells were fixed in 2% glutaraldehyde (Electron Microscopy Sciences) with 2.5% formaldehyde (Electron Microscopy Sciences) in 0.1 M sodium cacodylate buffer (pH 7.4) for 2 h, at room temperature, and post fixed in 1% osmium tetroxide (Electron Microscopy Sciences) diluted in 0.1 M sodium cacodylate buffer. Samples were then dehydrated and embedded in Epon resin (TAAB). Ultra-thin 50 nm sections were cut on an RMC Ultramicrotome (PowerTome) using Diatome diamond knives, mounted on 200-mesh copper grids (Electron Microscopy Sciences), and stained with uranyl acetate substitute (Electron Microscopy Sciences) and lead citrate (Electron Microscopy Sciences) for 5 min each. Sections were then examined under a JEOL JEM 1400 transmission electron microscope (JEOL) and images were digitally recorded using a CCD digital camera Orius 1100W.

### 2.22. Statistical Analysis

Two-way ANOVA followed by Tukey post hoc tests were used to test the effect of cell line and microenvironmental conditions on different biomarkers (HIF-1α, lactate, and ATP levels) and functional responses (invasion, proliferation, apoptosis). Differences were considered significant for *p*<0.05. All experiments were performed in triplicates and three replicates were conducted for each independent experiment. The results are presented as the average and standard deviation of these independent assays.

## 3. Results and Discussion

Oxygen and glucose deprivation are salient features of advanced stage bladder cancer, with negative implications in disease outcome. However, the precise role played by these microenvironmental features burdening poorly vascularized tumour regions remains insufficiently understood. Here, we have addressed the functional and molecular plasticity of bladder cancer cells under these conditions, envisaging to better understand the microenvironmental pressures governing bladder cancer. We also pursued preliminary observations implicating hypoxia in glycome remodelling in ways that favour disease progression and dissemination^12^. Emphasis was set on understanding the synergistic impact of low oxygen pressure and glucose in protein *O*-glycosylation, which remains an unexplored matter. We hypothesize such changes may provide the molecular rationale to identify more aggressive cancer cells capable of supporting severe microenvironmental stress and disease perpetuation.

### 3.1. Functional plasticity under hypoxia and low glucose

To better understand the molecular adaptability of bladder cancer cells to hypoxia and low glucose, we have grown four widely studied bladder cancer cell models (RT4, 5637, T24, HT1197) under low oxygen concentrations (0.1% O_2_) and reduced glucose levels (≤10%) to mimic microenvironmental conditions encountered by cells growing far apart from blood vessels. Cells in hypoxia responded rapidly to these changes, stabilizing the hypoxia biomarker HIF-1α, which became more pronounced in the absence of glucose, supporting HIF-1α pivotal role in adaptive responses to microenvironmental stress (**Figure 1A**). Notably, HIF-1α was also increased in cells grown at normal oxygen pressure but with very low glucose, reinforcing the existence of a non-canonical regulation mechanism for HIF-1α stabilization regardless of oxygen availability, as previously described for other types of cancer cells^18–20^. We then quantified intracellular and extracellular L(+)-lactate levels through the detection of reduced products of lactate dehydrogenase. Although differences were visualized according to the cell line, in general, lactate increased under low oxygen and was rapidly extruded to the extracellular space, suggesting the adoption of anaerobic glycolysis as main bioenergetic pathway and capacity to maintain intracellular homeostasis, as extensively supported in the literature^21,22^. Low glucose that mimics the Warburg effect also increased lactate as result of aerobic glycolysis^23,24^. However, lactate remained in the intracellular compartment, suggesting that low oxygen may be critical for activating extrusion mechanisms. On the other hand, the combined effect of hypoxia and low glucose reduced lactate close to vestigial levels, strongly supporting the activation of alternative energy producing pathway to glycolysis.

**Figure 1.**
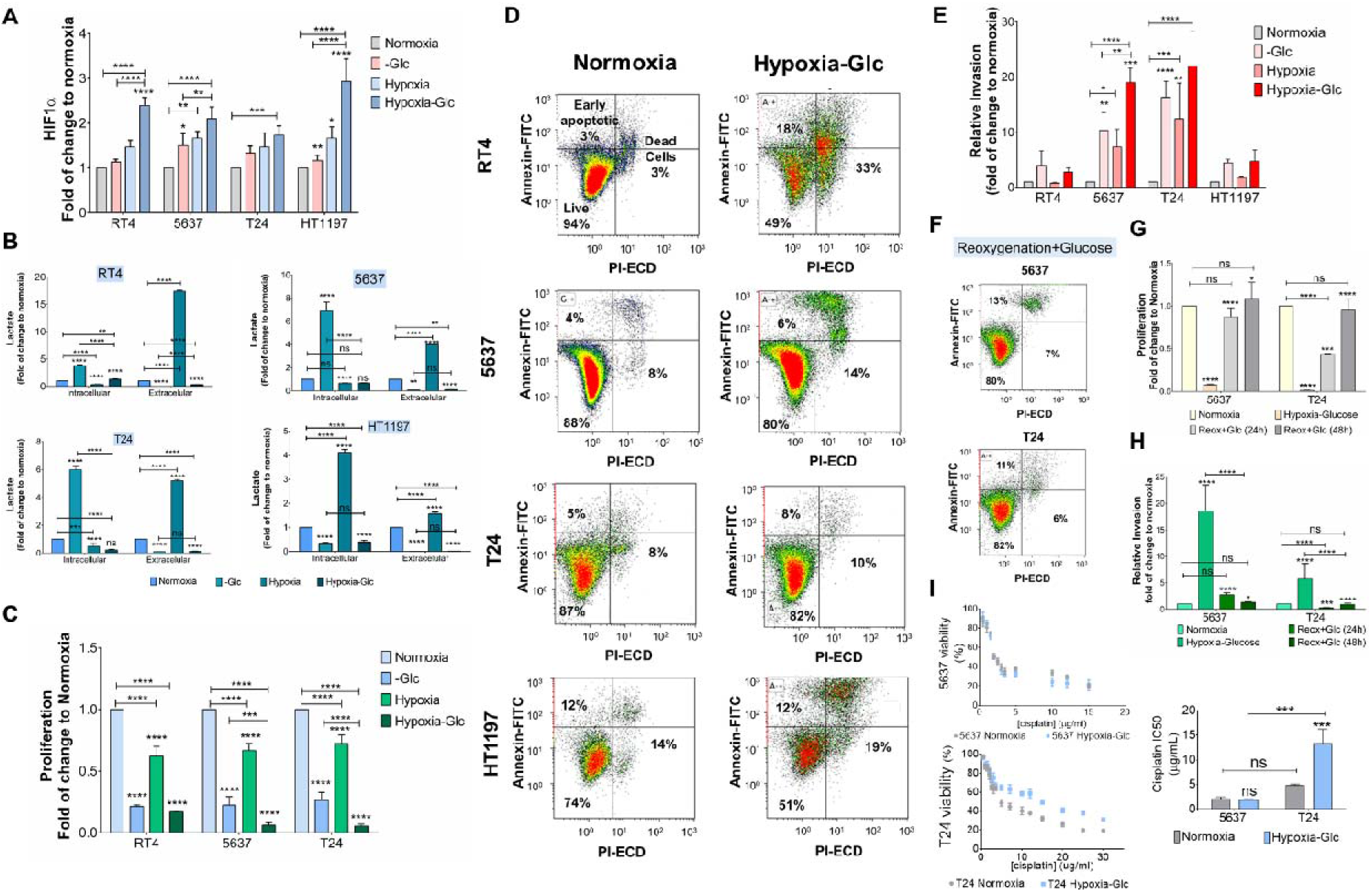
Bladder cancer cells tolerate well hypoxia and low glucose (herein termed Hypoxia-Glc) and adopt quasi-quiescent and invasive behaviour. **A) Hypoxia and low glucose significantly increase HIF-1**α **expression in bladder cancer cell lines.** HIF-1α is significantly stabilized either under low oxygen or glucose concentrations, irrespectively of the cell line. This effect becomes significantly more pronounced when the two microenvironmental stressors are combined. **B) Bladder cancer cells grown in hypoxia and low glucose produc residual levels of lactate.** Intracellular lactate increases in cells exposed to low glucose concentrations, whereas extracellular lactate increases in cells grown in hypoxia. The combination of both factors reduces lactate to vestigial levels**. C) Hypoxia and low glucose significantly decrease cell proliferation.** Individually, low oxygen or low glucose decrease bladder cancer cells proliferation capacity. Their combination significantly enhances this effect for all cell lines. **D) Bladder cancer cells tolerate well hypoxia and low glucose, maintaining cell viability.** Environmental stress resulting from the combined effect of hypoxia and low glucose did not impact significantly on the viability of 5637 and T24 cells. Both RT4 and HT1197 cell models reduced cell viability by 50% under these conditions, suggesting reduced capacity to adapt. **E) Bladder cancer cells become more invasive under hypoxi and low glucose** *in vitro*. The 5637 and T24 cell models showed more capacity to invade Matrigel *in vitro* when exposed to oxygen and glucose shortage separately. Moreover, invasion was significantly enhanced when the two stimuli were combined. The RT4 and HT1197 cell lines did not present significant invasive capacity in our study conditions. **F) Bladder cancer cells showed remarkable capacity to accommodate microenvironmental changes with minimal impact on cell viability.** Restoration of oxygen and glucose did not impact on cell viability, supporting high plasticity to accommodate drastic microenvironmental changes. **G) Bladder cancer cells restored basal proliferation after reoxygenation and glucose reconditioning.** After 24h of reoxygenation with restoration of glucose, 5637 and T24 cells significantly regained proliferative capacity. After 48h, cells regained basal proliferation, supporting the plasticity of these cells to endure microenvironmental challenges. **H) Bladder cancer cells reinstate basal invasion after reoxygenation and glucose restoration.** After 24h of reoxygenation with restoration of glucose, 5637 and T24 cells significantly reduce invasion. After 48h, their original invasive capacity was reinstalled. **I) Hypoxia and low glucose increased T24 cells resistance to cisplatin.** In normoxia, 5637 and T24 cells showed similar IC50 for cisplatin. In hypoxia and low glucose, T24 increased its tolerance to cisplatin over a wide range of concentrations, including its IC50, whereas 5637 remained unchanged. ns: not significative; \**p* < 0.05; \*\**p* < 0.01; \*\*\**p* < 0.001; **** *p* < 0.0001 (two-way ANOVA Tukey post hoc test)

Concomitantly to these molecular adaptations, we observed a striking decrease in cell proliferation (10-15-fold) in all cell lines under hypoxia, which was significantly enhanced upon glucose suppression (**Figure 1C**). Cell cycle arrest in S/G2 transition was later confirmed by flow cytometry (**Figure S1-Supporting Information**). Therefore, we hypothesized that cells may be arrested in late S phase due to depletion of the substrates required for DNA synthesis^25^, which was later confirmed by metabolomics. Interestingly, prolonged G2 arrest has been extensively described as a relevant therapy-escape mechanism after exposure to DNA damaging agents^26^, which was also later confirmed for these cells. Notably, viability of 5637 and T24 cells was not affected after 24 h of microenvironmental stress, as highlighted by little changes in the percentage of apoptotic and pre-apoptotic cells in comparison to normoxia (**Figure 1D**). In contrast, RT4 and HT1197 cells viability was decreased by ~50%. Strikingly, 5637 and T24 cells also responded to oxygen and glucose shortage by increasing invasion in Matrigel *in vitro* (**Figure 1E**). Also, the suppression of glucose impacted more on invasion than the removal of oxygen, highlighting the pivotal role played by this nutrient in cancer^27^. In parallel, we assessed the activity of matrix metalloproteinase-2 (MMP-2) and MMP-9, which are well known molecules supporting invasion in bladder cancer^28^. However, neither 5637 nor T24 cells exhibited changes in metalloproteinase activity under microenvironmental stress that could support the exuberant increase in invasion, strongly suggesting the adoption of alternative mechanisms (**Figure S2-Supporting Information**). Finally, reoxygenation and access to glucose significantly restored proliferation by 50% after 24 h and induced a massive drop in invasion without inducing apoptosis, suggesting little oxidative stress from drastic alterations in the microenvironment (**Figures 1F-H**). Collectively, this demonstrated that some subpopulations of bladder cancer cells are well capable of accommodating hypoxia induced stress, facing proliferation arrest supported by anaerobic metabolism, while concomitantly acquiring more aggressive and motile phenotypes. Moreover, it demonstrates that oxygen and glucose levels act as an on-off switch between proliferation and invasion. Interestingly, resistance to cancer cell death, early stop in proliferation, and activation of invasion traits have been closely linked to lactic acidosis as result of either hypoxia or glucose shortage^29^. However, our observations support the notion that bladder cancer cells may adopt similar behaviours in the absence of lactate in response to the combination of these microenvironmental factors. Finally, we addressed tolerance to cisplatin, generally used in the clinics against less proliferative tumour cells. Under hypoxia and glucose deprivation, bladder cancer cells either maintained or significantly increased tolerance to cisplatin as observed for T24 cells (**Figure 1I**). Collectively, we have portrayed the decisive role played by these microenvironment features in bladder cancer cells aggressiveness, setting the rationale for more in-depth molecular studies envisaging the identification of relevant molecular targets for precision medicine.

### 3.2. Bladder cancer cells transcriptome remodelling under hypoxia and low glucose

Hypoxia is known to induce significant transcriptome remodelling in cancer cells, allowing rapid adaptation to rising microenvironmental challenges. However, its combination with glucose deprivation remains poorly understood, as these events are generally studied separately. As such, we have performed comparative whole transcriptome analysis by RNA-Seq of 5637 and T24 cells under concomitant exposure to these conditions (**Figure 2**). According to the principal components analysis (PCA) in **Figure 2A**, the greatest variance is depicted by PC1 (95% variance), concerning differences between the cell lines. PC2 (4% of the variance) portrayed differences between normoxia and hypoxia with low glucose. Collectively, the two cell lines showed markedly different transcriptomes but also common responses to hypoxia and glucose shortage. Moreover, the global transcriptional change across the groups was visualized through a volcano plot (**Figure 2B**), which highlighted 3003 differentially expressed genes in hypoxia and glucose deprivation (1408 upregulated, 1595 downregulated), thus supporting significant transcriptome remodelling. A bi-clustering heatmap involving the top 30 differentially expressed genes sorted by their adjusted *p*-value also allowed to identify co-regulated genes across the different microenvironments (**Figure 2C**). Between the most differentially expressed and upregulated genes under hypoxia and glucose deprivation are *KRT17, GADD34/PPP1R15A, ETS1, DDIT4/REDD1, HK2, PFKFB3, DDIT3, SLC2A3, TAGLN, SLC22A15, SH3D21, DEPP1, RORA, ANGPTL4, ELOVL6*, *FOSB*, *LMNB1*, and *EGR1*. Together, these constitute a panel of biotic stress activated genes driving systemic changes at the transcriptomic level towards more undifferentiated (*KRT17*, *FOSB*)^30,31^, poorly proliferative (*LMNB1*)^32,33^, and less prone to programmed cell death and anoikis (*GADD34/PPP1R15A*, *DDIT4/REDD1*, *PFKFB3*, *ANGPTL4*) phenotypes^34–37^. Moreover, the negative regulation of suppressor factors (*EGR1)*^38^, allied to the promotion of immunosuppressor or tolerogenic (*GADD34/PPP1R15A, DDIT3)* programs^39–43^ leads to enhanced invasive/migratory (*ETS1*, *DDIT4/REDD1*, *TAGLN*) capacities^44–46^, as previously demonstrated. Finally, the pressing need for optimized energetic pathways facing nutrient shortage drives enhanced glucose uptake (*SLC2A3/GLUT3)*^47,48^ and lipid catabolism (*RORA, ANGPTL4)*^49,50^. The later includes the upregulation of carnitine transporters (*SLC22A15)*^51^, while counteracting lipogenesis (*ELOVL6)* through rate limiting enzymes downregulation^52^ and promotion of autophagic events (*DDIT4, HK2, PFKFB3, DEPP1*)^53–56^. Significantly differentially expressed genes were then clustered by gene ontology and the enrichment of gene ontology terms was tested using Fisher’s exact test (**Figure 2D**). Under hypoxia and glucose deprivation, cells fine-tune transcription of genes involved in cell-cell adhesion, cell proliferation, programmed cell death, DNA damage stimuli, metabolism reprogramming, and oxidation-reduction processes. Altogether, this denotes an intensive transcriptome remodelling towards adaptation to biotic stress and DNA damaging factors, supporting functional assays. Finally, to explore cellular processes and their dynamics, a functional interaction network was obtained using Cytoscape Software CluePedia and ClueGO plugins for single cluster analysis and comparison of gene clusters (**Figure 2E**). The most prominent groups of nodes include cellular responses to oxygen levels and biotic stimuli, as well as carbohydrate metabolism and inflammatory response. This suggests an underlying correlation between biotic stimuli, herein translated by oxygen and glucose shortage, and carbohydrate metabolism and biosynthesis, which could ultimately impact on complex networks as inflammatory responses. Moreover, there are suggestions that these cells may resemble with fat cells, suggesting the adoption of similar lipid metabolic pathways as suggested by *RORA* and *ANGPTL4* up-regulations (**Figure 2C**). Collectively, BC showed remarkable transcriptomic adaptability to microenvironmental stress. Transcriptomics analysis further supported all main functional alterations accompanying adaptation to low oxygen and glucose, providing the molecular foregrounds for their plasticity facing hostile conditions. Moreover, it strongly suggests the adoption of a lipolytic metabolism, which was latter assessed. Furthermore, despite the remarkable differences between the two cell lines, common molecular grounds were observed in terms of response to stress.

**Figure 2.**
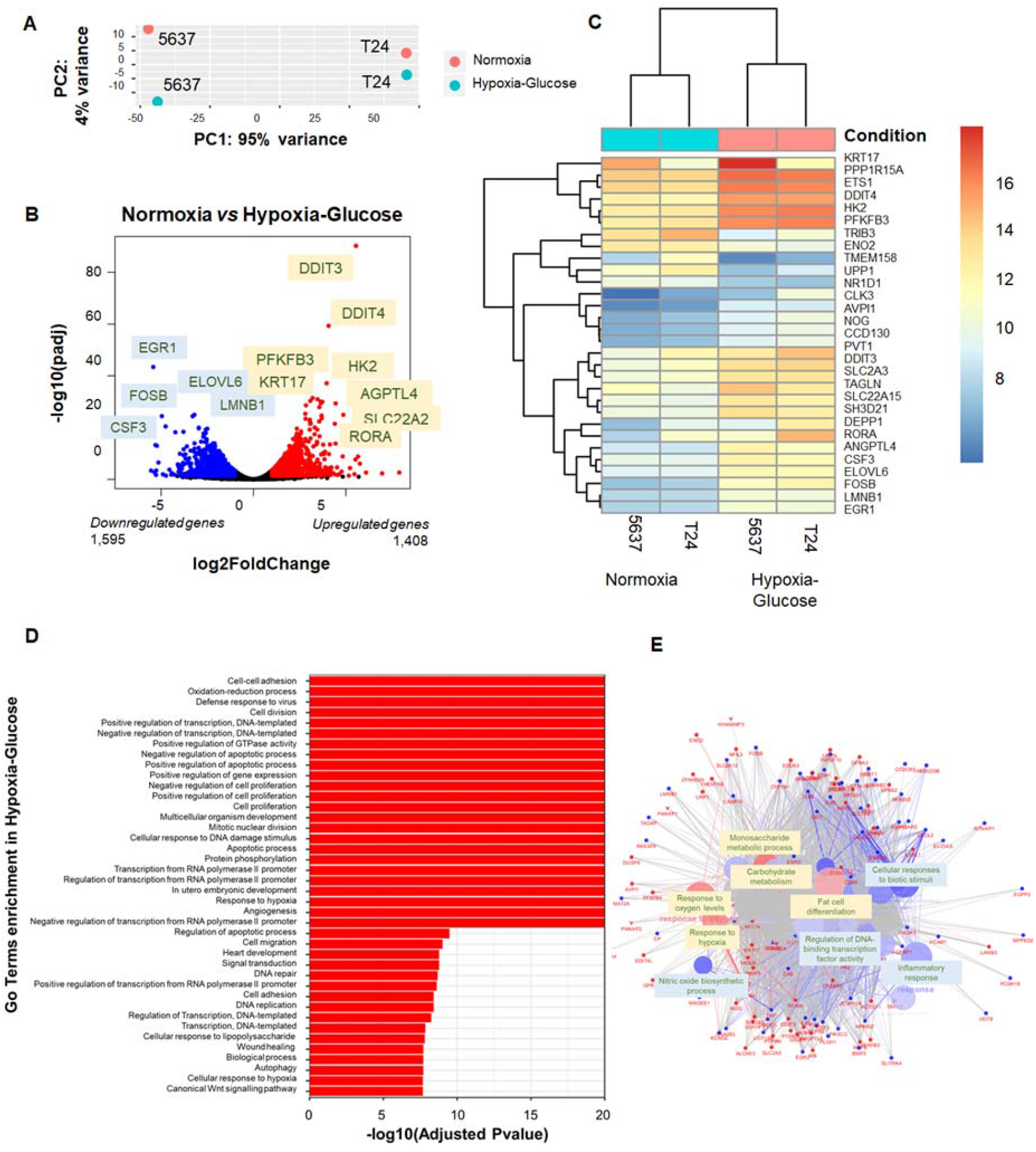
Bladder cancer cell lines under hypoxia and low glucose experience profound transcriptome remodelling, supporting the acquisition of more aggressive phenotypes. **A) Bladder cancer cells exhibit markedly different transcriptomes but also show common responses to hypoxia and low glucose.** Principle components analysis (PCA) of transcriptomics data showed that the greatest variance explained by PC1 (95% variance) concerns differences between the two cell lines. PC2, explaining 4% of the variance, showed marked differences between normoxia and hypoxia with low glucose, irrespectively of the cell line. PCA supports that, despite marked differences between cell lines, there are common responses to different microenvironmental features. **B) Volcano plot highlighting global transcriptional change between normoxia and hypoxia plus low glucose.** Exposure to hypoxia and glucose deprivation changed the expression of 3003 genes (1408 upregulated, 1595 downregulated), supporting significant transcriptome remodelling. **C) Bi-clustering heatmap concerning the top 30 differentially expressed genes showed co-regulations in hypoxia and low glucose that support proliferation arrest, resistance to cell death and invasion.** A bi-clustering heatmap was performed using the top 30 differentially expressed genes, sorted by their adjusted *p*-value, by plotting their log2 transformed expression values in samples. **D) Enrichment of gene ontology terms for differently expressed genes highlights alterations in main pathways associated with cell-cell adhesion**, **cell proliferation, and resistance to cell death. E) Functional interaction networks highlight several cellular processes activated in response to oxygen levels and biotic stimuli, including changes in carbohydrate metabolism and inflammatory responses.** Main functional nodes are highlighted. Only pathways with a *p* ≤0.001 were considered.

### 3.3. Metabolic adaptive responses to hypoxia and low glucose

To gain more insights on the metabolic reprogramming induced by hypoxia and glucose deprivation, we performed an untargeted metabolomics study by LC-MS/MS on 5637 and T24 cells. As highlighted by **Figures 3A-C**, T24 and 5637 cells present different metabolic fingerprints under normoxia, in line with distinct molecular backgrounds already observed at the transcriptome level (**Figure 2**). Nevertheless, BC cells presented similar metabolomic responses facing low oxygen and glucose, which were characterized by a statistically significant reduction in the levels of 85 metabolites and increments in 8 species associated with main cell pathways. A discriminant PLS-DA analysis also showed a clear separation between experimental conditions (**Figure 3C**), which in agreement with the volcano plot (**Figure 3A**) highlighted top metabolites contributing to group discrimination. Increased metabolites included 2-phenylaminoadenosine, xi-5-hydroxidecanoic acid and several fatty acid-carnitine derivatives, while uridine diphosphate glucose (UDP-glucose), uridine diphosphate N-acetylgalactosamine (UDP-GalNAc), citric acid, and glucoronic acid levels were substantially decreased. Particularly, the generation of adenosine and adenosine receptors agonists such as 2-phenylaminoadenosine (A2 selective ligand) have been found increased under hypoxic conditions, exerting immune regulatory functions^57^. The engagement of adenosine A2 receptors frequently leads to immunosuppressive pathways, including inhibition of cytotoxicity and secretion of pro-inflammatory cytokines promoted by activated immune cells^58^. A potential role in angiogenesis promotion has also been suggested by several authors^58^. On the other hand, 5-hydroxydecanoate (5-HD), a specific mitochondrial ATP-sensitive K^+^ channel inhibitor, has been described to attenuate the loss of mitochondrial transmembrane potential, the increase in the formation of reactive oxygen species, and proteasome inhibitor-induced apoptosis by suppressing the activation of caspase-8 and Bid-dependent pathways^59^. Finally, highly exacerbated levels of fatty acid-carnitine derivatives (pentadecanoylcarninite, L-palmitoylcarnitine, heptadecanoylcarnitine, stearoylcarnitine, arachidylcarnitine; **Figures 3A-C**) highlight active translocation of long-chain fatty acids across the inner mitochondrial membrane for subsequent *β*-oxidation, which was also suggested by the highly lipolytic transcriptomic profile of hypoxic BC cells^60^. Similar observations have been made by other authors in the urine of advanced stage patients^61,62^, reinforcing the close link between this metabolic phenotype and aggressiveness. Hypoxic and glucose deprived cells also significantly reduced the levels of citric acid, which might result from the combined effect of loss of mitochondria due to mitophagy and reduced tricarboxylic acid (TCA) cycle activity^63^. Moreover, in the event of a reduction in mitochondrial citrate production, other pathways could supply cytosolic citrate, including the reversed isocitrate dehydrogenase (IDH) reaction; nevertheless, *IDH1* was also found downregulated under these experimental conditions, further reinforcing citric acid overall reduction. Furthermore, reduced concentration of citrate in cancer cells has been described to favour resistance to apoptosis and cellular dedifferentiation^63^, thus in agreement with functional and transcriptomics studies. The significant reduction in UDP-GalNAc, a key sugar nucleotide for the initiation of protein *O*-glycosylation, supports major alterations in the glycophenotype of cancer cells. This may be intimately related with decreased glucose metabolism via the pentose phosphate pathway (PPP) as result of low glucose availability, which is essential for nucleotide sugar biosynthesis, including UDP^64^. Glucose shortage may also compromise anabolic processes such as carbohydrate synthesis^65^, further contributing to low UDP-GalNAc. Finally, glucose shortage may also explain the low levels of glucuronic acid that directly derives from its oxidation^66^ and likely have a profound impact on glycosaminoglycans and proteoglycans biosynthesis, which are key extracellular molecules involved in an onset of oncogenic events^11,67^. Moreover, an integrated enrichment overview has evidenced carnitine biosynthesis and carnitine precursors degradation, like lysine and methionine, as mainly enriched pathways (**Figure 3 D and E**). Fatty acid metabolism and mitochondrial *β*-oxidation are also prominent, in agreement with the lipolytic phenotype highlighted by transcriptomic analysis (**Figure 3D**). Collectively, our findings support that hypoxia and low glucose induce lipid catabolism as the main bioenergetic pathway. Moreover, according to several studies, the metabolites produced under hypoxia and low glucose are directly linked to poorly immunogenic and undifferentiated phenotypes and more advanced stages of the disease^68,69^.

**Figure 3.**
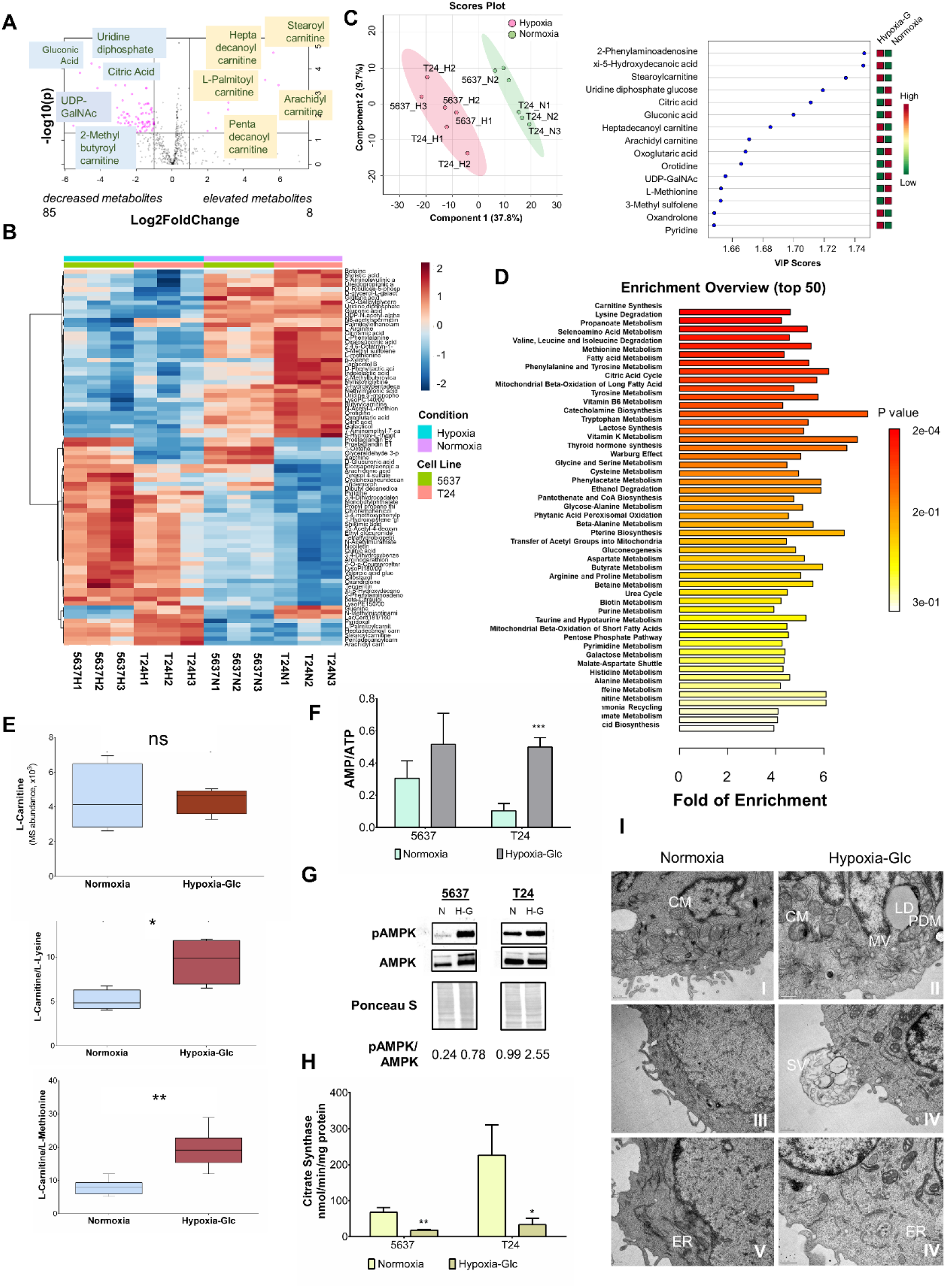
Hypoxia and low glucose induce a shift from glycolytic to lipolytic metabolism, accompanied by a significant reduction in the number of intact mitochondria by autophagy. **A) Volcano plot highlighting major metabolome alterations in response to hypoxia and low glucose.** Eighty-five metabolites were downregulated, namely uridine diphosphate, relevant for glycogenesis and the synthesis of several nucleotide sugars such as UDP-GalNAc, and glucuronic acid, which together support significant changes in the glycome of cancer cells. Citric acid was also underrepresented, supporting the inactivation of the tricarboxylic acid cycle. On the other hand, 8 metabolites corresponding to carnitine long-chain fatty acids esters were increased, suggesting active fatty acids transport to mitochondria for subsequent *β*-oxidation. **B) Discriminating heat map and C) PLS-DA analysis revealed similar metabolic responses by 5637 and T24 cells under microenvironmental stress.** The 5637 and T24 cells showed similar metabolomes under hypoxia and low glucose, denoting analogous metabolic responses facing microenvironmental stress. **D) Pathway enrichment analysis supports major alterations in the metabolome at different levels, including the adoption of fatty acid** β**-oxidation as main bioenergetic pathway in nutrient deprived bladder cancer cells.** A wide array of key metabolic pathways was influenced by deprivation of oxygen and glucose, with emphasis on carnitine biosynthesis and carnitine precursors degradation, as lysine and methionine, supporting fatty acid *β*-oxidation. **E) Hypoxia and low glucose induced lysine and methionine degradation to support carnitine biosynthesis.** Under hypoxia and low glucose carnitine levels were maintained at the expenses of L-lysine and L-methionine degradation, supporting downstream lipids *β*-oxidation. **F) AMP/ATP ratio increases under hypoxia and low glucose accompanied by G) activation by phosphorylation of 5-AMP-activated protein kinase (AMPK).** Increased AMP/ATP ratios were observed due to increased AMP and decreased ATP, supporting impaired oxidative phosphorylation. In addition, an increase in pAMPK/AMPK ratios, mostly explained by increased pAMPK, was observed under microenvironmental stress, consistent with adoption of catabolic processes and potentially mitophagy. **H) Citrate synthase activity is decreased under hypoxia and low glucose.** Citrate synthase activity is significantly decreased in cells under hypoxia and low glucose, suggesting a reduction in the number of matrix intact mitochondria. **I) Hypoxia and low glucose induce major morphological changes in cancer cells, including mitophagy events.** Bladder cancer cells morphological aspects were inquired by TEM. Panel I shows normal cytoplasm mitochondria (CM) morphology in normoxia. Panel II demonstrates a lower number of mitochondria of apparently compromised nature in hypoxia and low glucose. Signs of mitophagy were evident. Lipid droplets (LD), peridroplet mitochondria (PDM), and multicellular vesicles (MV) were also clearly observable. Panel III reveals the basal membrane activity of bladder cancer cells. Panel IV highlights higher membrane activity under stress, translated by pronounced shedding of vesicles (SV). Panel V displays typically long endoplasmic reticulum (ER) sections in normoxia. Panel VI shows short and disorganized ER cisternae under stress, suggesting prominent disorganization of secretory pathways. ns: not significative; \**p* < 0.05; \*\**p* < 0.01; \*\*\**p* < 0.001 (Student’s T test for panel E; two-way ANOVA Tukey post hoc test for Figures E-G)

Furthermore, we observed an increase in AMP/ATP ratios **(Figure 3F)** accompanied by the activation by phosphorylation of 5-AMP-activated protein kinase (AMPK; **Figure 3G**) under hypoxia and low glucose, as expected for cells under microenvironmental stress^70,71^. Increased AMP/ATP ratio was governed by a major decrease in ATP levels, reinforcing the hypothesis of impaired oxidative phosphorylation. In parallel, increased pAMPK/AMPK ratios, mostly driven by higher pAMPK, have been shown, as previously observed in more aggressive bladder tumors^72^. These findings were consistent with adoption of catabolic processes such as fatty acid *β*-oxidation and potentially mitophagy^73^, which was later reinforced by the decrease in citrate synthase activity **(Figure 3H)** and TEM **(Figure 3I)**. In fact, TEM analysis evidenced a drastic decrease in the number of intact mitochondria, allied to evident mitophagy events (**Figure 3I-I and II**), translated by outer mitochondrial membrane-associated degradation and matrix sectioning. Lipid droplet (LD)-associated mitochondria, also known as peridroplet mitochondria (PDM), were also observed under hypoxic conditions **(Figure 3I-II)**, suggesting the existence of metabolically distinct subpopulations within the individual cell involved simultaneously in fatty acid oxidation and LD formation^74^. Under microenvironmental stress, vesicles shedding was evident **(Figure 3I-III and IV)**, highlighting cellular communication events in response to microenvironmental cues that should be carefully investigated in future studies. Finally, stressed cells showed considerably short and disorganized endoplasmic reticulum (ER) cisternae, contrasting with typically longer ER sections presented by cancer cells in normoxia **(Figure 3I-V-VI)**, demonstrating prominent disorganization of secretory pathways and potentially protein *O*-glycosylation changes^75^. Overall, biotic stress potentiated catabolic processes characterized by fatty acid oxidation and mitophagy as well as reduced anabolic processes that support carbohydrate biosynthesis and cellular proliferation. Moreover, it promoted a generalized disorganization of secretory organelles, which may also contribute to global glycosylation alterations.

### 3.4. Transcriptomics and Metabolomics Joint Pathway Analysis

A joint pathway analysis, combining transcriptomics and metabolomics, was then pursued to gain more insights on the plasticity of BC cells. We found changes in glycolysis and glyconeogenesis as well as glycerolipids metabolism, which support the adoption of a lipolytic rather than a glycolytic metabolism. In addition, we observed major inhibition of different pathways that may directly or indirectly impact on cell glycosylation patterns by impairing nucleotide or sugars biosynthesis. Namely, inhibition of fructose and mannose metabolism was observed, both producing sugar intermediates necessary to fuel the hexosamine biosynthetic pathway (HBP) and others leading to nucleotide sugar production, including UDP biosynthesis^76–78^, which constitutes a major limitation for building mature glycans. In agreement with these observations, mucin-type *O*-glycans biosynthesis emerged as one of the most altered pathways, supporting previous observations (**Figure 4A**). This was driven by significant reduction in UDP-GalNAc levels under hypoxia and low glucose in both cell lines **(Figure 4B)**. Interestingly, no changes in gene expression were observed for key enzymes involved in the hexosamine biosynthetic pathway, namely *MPI*, *GNE*, *GALE*, encoding for mannose-6-phosphate isomerase, UDP-GlcNAc-2-epimerase/ManAc kinase and UDP-glucose 4-epimerase/UDP-galactose 4-epimerase, respectively (data not shown). However, several glycogenes encoding different polypeptide *N*-acetylgalactosaminyltransferases responsible by the initial step of protein *O*-glycosylation (*GALNT1*, *GALNT3*, *GALNT7*, *GALNT10*, *GALNT12;* **Figure 4C**) were downregulated in hypoxia and low glucose, suggesting that microenvironmental stress could directly impact on the number of protein *O*-glycosites. We also observed a downregulation of *C1GALT1C1/COSMC*, encoding for C1GALT1-specific chaperone 1. This chaperone is essential for core 1 β1-3-galactosyltransferase 1 (T synthase) function and determines cell capacity to elongate glycans beyond the initial Tn structure. Moreover, its abrogation has been associated with malignancy and worst prognosis in different cancers^79,80^. Collectively, these observations strongly suggest that hypoxia and glucose shortage act as microenvironmental pressures towards the biosynthesis of immature truncated *O*-glycans and, potentially, less densely glycosylated proteins. Remarkably, we have previously demonstrated that bladder tumours present significant alterations in their glycosylation, translated by the expression of shorter glycans, with major implications in bladder cancer progression and dissemination^12,13,81–83^. Nevertheless, the underlying microenvironmental features driving these events remained so far undisclosed.

**Figure 4.**
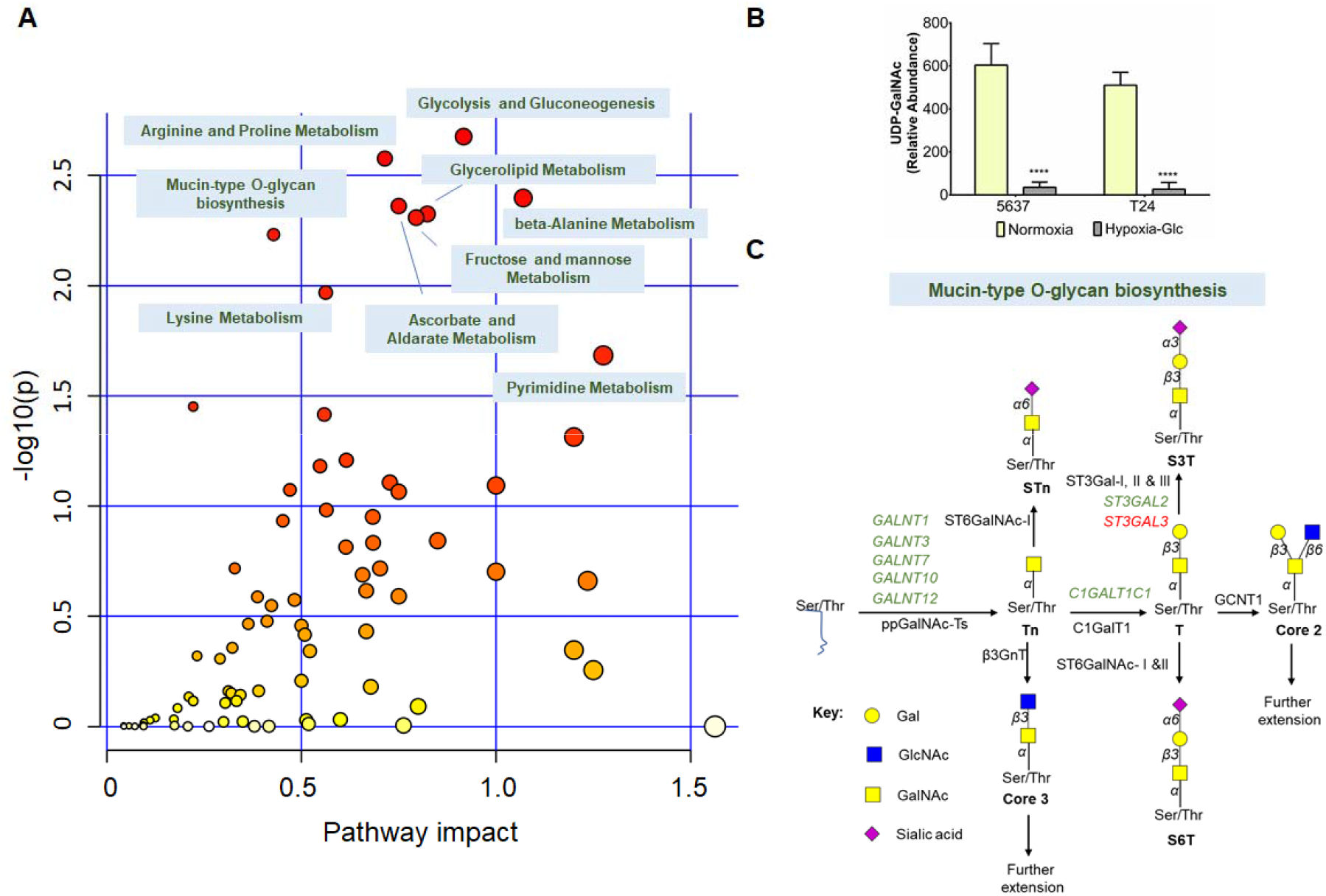
**A) Joint metabolome and transcriptome pathway analysis supports major remodelling of bladder cancer cells metabolic pathways accompanying adaptation to hypoxia and low glucose.** Joint pathway analysis supports changes from glycolytic to lipolytic metabolism and inhibition of relevant pathways supporting nucleotides and sugars biosynthesis, ultimately negatively impacting on *O*-GalNAc glycans biosynthesis. **B) Bladder cancer cells under microenvironmental stress exhibited significantly decreased UDP-GalNAc levels in comparison to normoxia.** \*\*\**p* < 0.001 (two-way ANOVA Tukey post hoc test for panels F and H) **C) Schematic representation of mucin-type *O*-glycan biosynthesis highlighting main transcriptomic changes in relevant biosynthetic enzymes.** Bladder cancer cells displayed significant downregulation (highlighted in green) of several *GALNT*s involved in *O*-glycans initiation and *C1GALT1C1*, which encodes for a key chaperone of C1GalT1 that drives glycans elongation. ST3GAL3 was the only upregulated glycogene in this pathway (highlighted in red), suggesting increased *O*-3 Gal sialylation.

### 3.5. *O*-glycomics

Glycomics analysis was conducted to assess the hypothesis of *O*-glycans biosynthesis antagonization by the microenvironment. This was performed exploiting the Tn mimetic benzyl-α-GalNAc as a scaffold for further *O*-chain elongation **(Figure 5)**. Cells were also reoxygenated and glucose levels were restored to evaluate *O*-glycome plasticity. According to **Figure 5A**, low oxygen and glucose significantly reduced the amount of extended *O*-glycans (herein defined as glycans resulting from modification of the Tn antigen) produced by both cell lines in comparison to normoxia, as suggested by joint pathway analysis. This effect was mostly driven by the removal of glucose, translating the key importance of this metabolite for *O*-glycosylation pathways. Hypoxia enhanced the net effect induced by glucose, as clearly highlighted by **Figures 5B-E**. More detailed glycomic characterization in **Figure 5B** showed that both cell lines abundantly express fucosylated (*m/z* 746.40; type 3 H-antigen) and sialylated T (*m/z* 933.48) antigens, also exhibiting several extended core 2 *O*-glycans of variable lengths, degrees of fucosylation and sialylation. Low amounts of shorter *O*-glycans such as core 3 (*m/z* 613.33) and STn (*m/z* 729.38) antigens could also be observed. However, the drastic reduction of oxygen and glucose tremendously impacted the glycome of cells, inducing a simple cell glycophenotype characterized by an accumulation of few short-chain *O*-glycans without chain extension beyond core 1 **(Figure B)**. The most abundant glycoform was core 3 and, to less extent, mono- (*m/z* 933.48) and di-sialylated (*m/z* 1294.65) T antigens (**Figures 5B-E**). T antigen fucosylation was almost completely inhibited under these conditions. Trace amounts of STn antigen could also be detected. Notably, core 3 expression is being reported for the first time in bladder cancer cells, being typical of the colorectal epithelium, where it plays a key role in homeostasis^84^. Interestingly, no extension of core 3 was observed, reinforcing the inexorable expression of shorter structures by these cells. Moreover, while the relative abundance of core 3 increases in relation to other glycans, its total amount remains mostly unchanged from normoxia to hypoxia with low glucose, as highlighted by **Figure 5C**. On the other hand, cells significantly reduce the total amount of core 1 structures, namely sialyl-T antigens. Collectively, these findings support reduced capacity to extend glycans to core 1, as previously suggested by the decreased expression of *C1GALT1C1,* encoding for C1GalT1-specific chaperone 1 (Cosmc**; Figure 4C)**. This likely results in an accumulation of immature Tn glycans, while maintaining core 3 biosynthesis steady, later confirmed by immunoassays (**Figures 5F-G)**. Finally, **Figure 5E** also clearly highlights that the inhibition of *O*-glycans extension beyond core 1 is mainly driven by the reduction in glucose, since some extended structures could still be observed under hypoxia. Moreover, bladder cancer cells regain the capacity to extend glycans after reoxygenation and reintroduction of glucose (**Figure 5E**), reinforcing the hypothesis of an on-off switch for *O*-glycosylation depending on the microenvironment.

**Figure 5.**
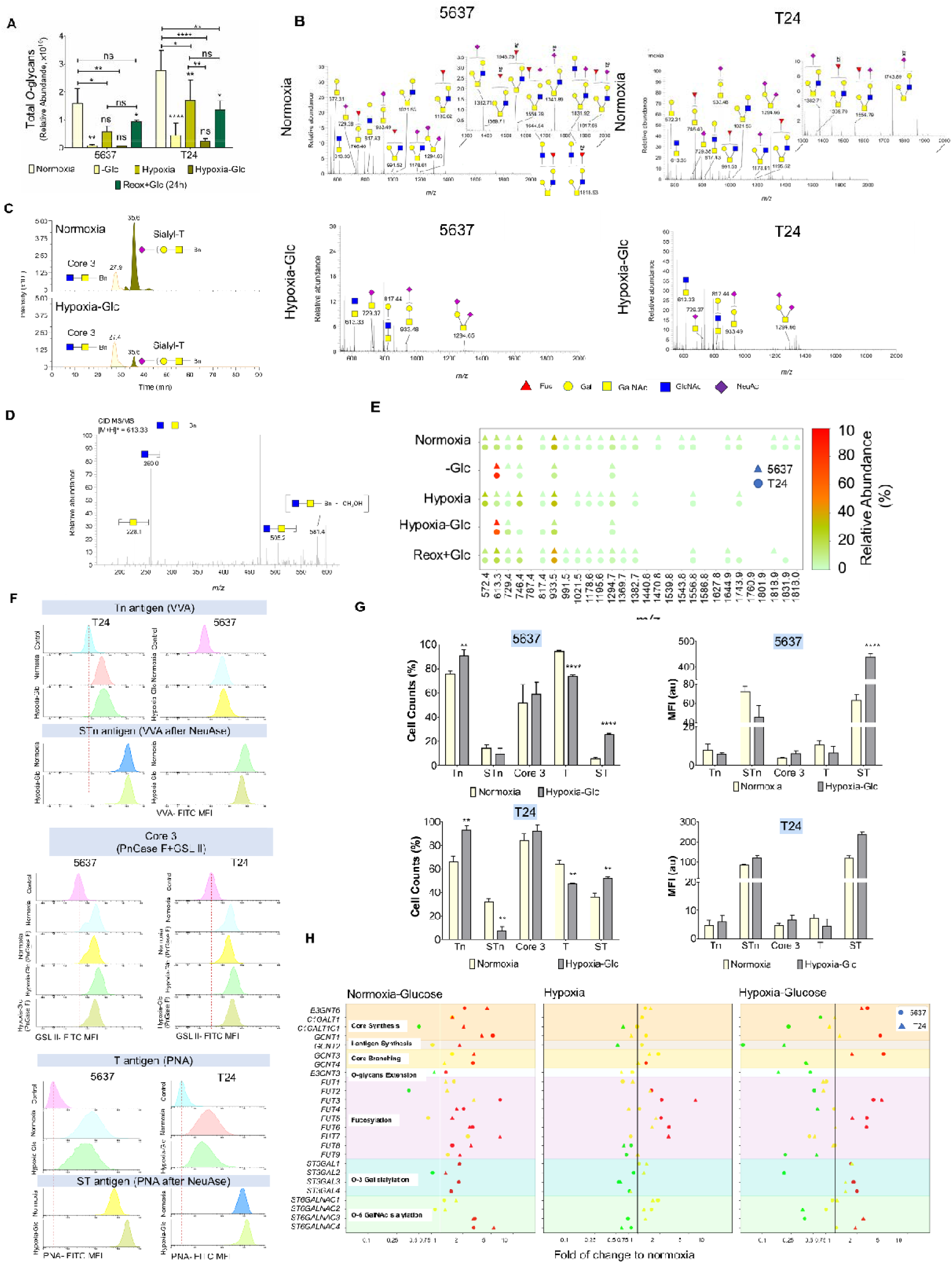
Hypoxia and low glucose antagonize *O*-glycans extension, inducing a simple cancer cell glycophenotype. **A) Hypoxia and low glucose induce a massive reduction in bladder cancer cells capacity t produce *O*-glycans.** Bladder cancer cells showed significantly decreased capacity to generate *O*-glycans under microenvironmental stress, mainly driven by the decrease in glucose availability. The combination of hypoxia and low glucose further enhances this effect, whereas reoxygenation and reposition of glucose restore normal glycosylation. Both cell lines responded similarly to microenvironmental stress. **B) Bladder cancer cells under hypoxia and low glucose present less abundant, simpler, and shorter glycomes, lacking extension beyond core 1 structures.** The full MS spectra highlight the loss of glycan chains extended beyond core 1 in cells exposed to hypoxia and low glucose. The absence/trace amounts of fucosylated core 1 were also noticeable in cells under microenvironmental pressure. Overall, under stress, cells acquired a simple cell glycophenotype, translated in this analysis by the presence of core 3 and sialylated T antigens. **C) Decrease in glycans abundance accompanying adaptation to the microenvironment is driven by the loss of extended structures as well as sialylated T antigens.** Extraction ion chromatograms (EIC) highlight a clear loss of sialylated T antigens in cells under stress. Notably, the abundance of core 3 remains constant. D**) Typical nanoLC-MS/MS spectrum for core 3.** Core 3 was identified for the first time in bladder cancer cell lines and the MS/MS highlights typical diagnostic fragment ions that confirm the structure. **E) Glycome heatmap highlighting the significant and reversible impact of glucose levels and its combination with hypoxia in the antagonization of** *O***-glycans extension.** Glycomics analysis show common responses in 5637 and T24 cells under different microenvironments, including the inhibition of *O*-glycans extension beyond core 1 under glucose deprivation. The reinstitution of glycan extension upon reoxygenation and reintroduction of glucose is also shown. **F and G) Lectin flow cytometry confirms a simple cell glycophenotype characterized by Tn/STn antigens and T and sialylated T in some subpopulations of cancer cells.** The Tn antigen was assessed with the VVA lectin. The STn antigen contribution arises from the comparison of VVA signals before and after sialidase (NeuAse) digestion. Core 3 was assessed through comparison of GLS II signals prior and after PNGaseF digestion. T antigens were estimated with PNA and sialylated T antigens with PNA after sialidase digestion. Except for an increase in sialylated T antigens, other glycans did not vary their expression levels from normoxia to hypoxia with low glucose. However, all cells acquired capacity to express the Tn antigen under stress. The percentage of cells expressing sialylated T antigens also increased, accompanying a decrease in the number of T antigen expressing cells, supporting higher sialylation. In summary, the Tn antigen became prevalent amongst cells under stress. Some subpopulations also showed capacity to express core 3, STn, and particularly the ST antigen. **H) Differential expression of glycogenes involved in glycan biosynthesis under low glucose, hypoxia, and the combination of both compared to normoxia.** Suppression of glucose is responsible by main alterations in glycogenes expression in comparison to hypoxia. Namely, glucose suppression induced significant glycogenes regulation involved in core 1 (*C1GALT1)*, 2 (*GCNT1),* and 3 (*B3GNT6*) synthesis, as well as core branching enzymes, and several fucosyltransferases and sialyltransferases. In hypoxia, glycogenes experience either mild downregulation or maintain their relative levels. The combination of both microenvironmental factors significantly enhanced major glycogenes downregulation, with emphasis on C*1GALT1* and *C1GALT1C1*, which dictate *O*-glycans extension (downregulation: green; no variation: yellow; upregulation: red). ns: not significative; \**p* < 0.05; \*\**p* < 0.01; \*\*\**p* < 0.001; **** *p* < 0.0001 (two-way ANOVA Tukey post hoc test).

To support these observations, we then assessed the cell surface glycome by flow cytometry, combining lectins and enzymatic digestions to expose glycans of interest. The Tn antigen was determined using the VVA lectin, whereas the STn antigen was assessed using the same lectin after sialidase digestion. The results were validated using B72-3 monoclonal antibody staining, which retrieved similar results (data not shown). Core 3 was indirectly assessed using the GSL II lectin, targeting GlcNAc residues at the nonreducing end of glycan chains. This was done to overcome the lack of commercially available antibodies for this glycan. To minimize possible cross-reactivity with *N*-glycans, GSL II was determined after PNGaseF digestion. Finally, the T antigen was characterized with the PNA lectin and sialylated T antigens determined by comparing the affinity of the lectin before and after neuraminidase digestion. According to **Figures 5F** and **G**, both cell lines expressed short-chain *O*-glycans, in accordance with mass spectrometry analysis. Moreover, hypoxia and glucose deprivation significantly increased the number of cells expressing the Tn and ST antigens, despite the inhibitory impact of this condition in *O*-glycosylation pathways. However, while the Tn antigen was detected in all cells, sialylated T antigens were found in less than half; yet with higher expression than in normoxia. Interestingly, the increase in the number of cells expressing sialylated T antigens was accompanied by a similar decrease in T antigen-expressing cells, suggesting higher sialylation in certain subpopulations. Moreover, there were little changes in the percentage of cells potentially expressing core 3 as well as in its intensity, as previously suggested by glycomics. The STn antigen levels were maintained constant. Taken together with mass spectrometry analysis, these findings support a massive stop in *O*-glycans extension, resulting in the accumulation of immature glycans such as the Tn/STn antigens and the maintenance of low levels of core 3. It also supports the existence of some subpopulations showing capacity to form more extended T and sialyl-T antigens, denoting significant glycome microheterogeneity. Moreover, it highlights the plasticity of *O*-glycosylation pathways in response to oxygen and glucose shortage.

Finally, we focused on understanding how the microenvironment influenced glycogenes expression. As such, we have re-analysed individually by RT-PCR the expression of a wide array of glycogenes potentially involved in *O*-glycans biosynthesis **(Figure 5H)**. Hypoxia promoted little alterations on glycogenes expression in comparison to conditions with low glucose. Nevertheless, we observed a decrease in *C1GALT1C1* in both cell lines, which was statistically significant for T24 cells, and may decisively contribute to inhibit *O*-glycans extension towards T antigen synthesis. On the other hand, the drastic reduction in glucose under normoxia upregulated *B3GNT6*, *C1GALT1*, and *GCNT1* genes, involved in core 3, 1, and 2 synthesis, respectively, as well as core branching enzymes, and several fucosyltransferases and sialyltransferases. Notably, *C1GALT1C1* expression was decreased in 5637 but increased in T24 cells. When hypoxia is associated with reduced glucose, there is a striking reduction in glycogenes expression (19/25 glycogenes), with emphasis on the downregulation of *C1GALT1* and *C1GALT1C1* in both cell lines, in accordance with whole transcriptome characterization **(Figure 2)**. Also, there was a significant overexpression of *B3GNT6* promoted by glucose suppression, which may contribute to sustain core 3 biosynthesis. In addition, we observed that hypoxia and glucose suppression downregulated *FUT1* and *FUT2*, which may explain the trace amounts of fucosylation of the T antigen. In addition, we observed an upregulation of several sialyltransferases that may contribute to ST overexpression (*ST3GAL1*, *ST3GAL3*, *ST3GAL4*), including ST3GAL3 that was previously observed by whole transcriptome analysis. Nevertheless, hypothesizing that the antagonization of *O*-glycans extension could be related to *C1GALT1* and *C1GALT1C1/COSMC* downregulation we further evaluated the levels of both proteins by western blot, which showed no changes with different microenvironmental stimuli **(Supplementary Figure S3).** Based on these findings, we hypothesize that other factors such as nucleotide sugars shortage and a net disorganization of organelles involved in the secretory pathway play a decisive role in this process.

In summary, we have demonstrated that hypoxia and low glucose induce a simple cell glycophenotype in bladder cancer cells, characterized by the absence of extended *O*-glycans, Tn/STn antigen expressions and, in some subpopulations of cancer cells, also T/ST. Moreover, we have demonstrated that glucose levels have major impact on the establishment of these glycophenotypes, which are exacerbated by hypoxia. Notably, our studies have linked these short forms of *O*-glycosylation to more aggressive forms of bladder cancer and poor prognosis^81–83^. In human tumours, we found these glycoforms in clustered cells, frequently in less proliferative tumour areas showing high HIF-1α levels, supporting close associations with microenvironmental niches^12,81^.

### 3.6. Simple cell glycoengineered models

We then developed a library of bladder cancer cells displaying different simple glycophenotypes presented by hypoxic and glucose deprived cells with the objective of gaining more knowledge on the biological role played by altered glycosylation in bladder cancer. Therefore, T24 cell line has been glycoengineered to hamper *O*-glycan extension beyond core 1 antigen. Accordingly, *C1GALT1* and *GCNT1* KO were produced using validated gRNAs^85^ through CRISPR-Cas9 technology. To increase STn antigen expression, human *ST6GALNAC1* has been knocked-in in *C1GALT1* KO cells. Briefly, three *C1GALT1* and GCNT*1* KO clones were selected according to their distinct indel profile, as determined by IDAA (**Supplementary Figures S4-5**). Sanger sequencing allowed detecting mutation sites in at least two different coding alleles (**Figure S4-5**). Further model validation was based on immunocytochemistry, flow cytometry, and orthogonal validation by MS (**Figure 6; Supplementary Figures S7-9**). Collectively, all controls displayed similar glycomes, characterized by high sialyl-T expression and vestigial/low Tn, STn, core 3, T antigens **(Figure 6)**, and extended core 2 structures **(Supplementary Figures S7-9)**, thus like wild type cells **(Figure 5)**. *C1GALT1* KO models were invariably characterized by a marked increase in Tn antigen **(Figure 6A; Supplementary Figure S6)**, with minor changes in STn and core 3 expressions. As expected, synthesis of T antigen and extensions beyond it were also not observed **(Figure 6A)**, confirming successful abrogation of C1GalT1 activity. By inducing *ST6GALNAC1* overexpression we were able to significantly increase STn at the expenses of the Tn antigen, while maintaining low core 3 **(Figure 6B; Supplementary Figure S7)**. Finally, *GCNT1* KO models mostly resulted in the expression of sialylated T antigens and complete loss of core 2 and other extended glycans. However, according to mass spectrometry **(Supplementary Figure S7),** it still presented high levels of core fucose, which are completely lost when cells are exposed to low oxygen and glucose. Efforts are ongoing to generate models lacking this type of fucosylation, to fully infer the impact of this poorly understood glycosignature in cancer. Overall, these models portrait the structural diversity of the glycome associated with hypoxia and glucose limitation and provided decisive tools to characterize its functional implications for disease.

**Figure 6.**
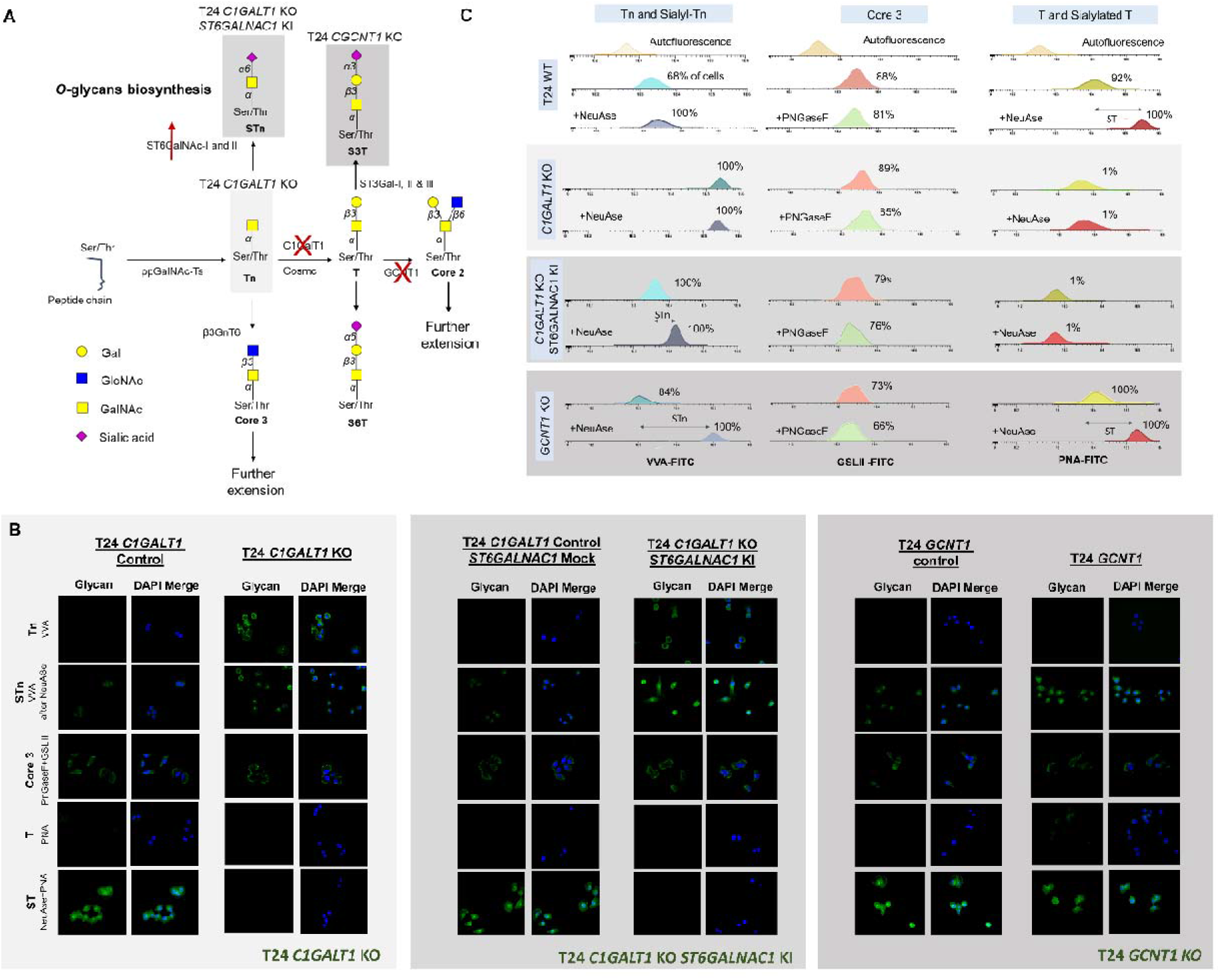
Library of simple cell glycoengineered models expressing glycophenotypes observed in hypoxic and glycose deprived cells. **A) Schematic representation of protein *O*-glycosylation pathways highlighting targeted glycogenes and associated glycophenotypes.** Light grey highlights the main glycan generated by *C1GALT1* KO; Medium grey highlights the main glycan generated by *C1GALT1* KO/*ST6GALNAC1* KI; Dark grey highlights the main glycan generated by *GCNT1* KO. **B and C) Glycoengineered T24 cell lines reflected the immature glycophenotype presented by cells exposed to hypoxia and low glucose.** T24 *C1GALT1* KO cells expressed highly immature Tn, little STn and no T or sialylated T antigens. Low core 3 expression was also observed. Stable *ST6GALNAC1* expression originating *C1GALT1* KO/*ST6GALNAC1* KI significantly increased STn, reducing Tn antigen levels. T24 *GCNT1* KO originated cells expressing high ST and, to less extent, STn antigens. No Tn was detected and vestigial amounts of core 3 were present.

### 3.7. Glycoengineered Cells Functional Assays

Glycoengineered cell models were used to interrogate the role played by glycosylation in decisive aspects of the disease, namely capacity to proliferate, invade, grow without anchorage, potentially metastasise, and tolerate chemotherapy agents. According to functional studies *in vitro*, *C1GALT1* KO cells were less proliferative than controls in both normoxia and hypoxia with limited glucose **(Figure 7A)**. However, Tn overexpression had no impact on invasion, which was mostly driven by changes in the microenvironment. Interestingly, *C1GALT1* KO cells presented higher capacity to grow in an anchorage-independent manner and resist anoikis, suggesting increased metastatic potential **(Figure 7A)**. Moreover, this model was more resistant to cisplatin over a wide array of drug concentrations and, consequently, presented a significantly higher IC50 than control **(Figure 7A)**. Finally, we explored the *in ovo* chick CAM model to assess the biological role of glycans *in vivo*. Notably, T24 CAM models have been previously developed and proven to reflect many histological and molecular features of human bladder tumours^86^, providing relevant platforms for functional studies. *C1GALT1* KO CAMs exhibited significantly smaller tumours when compared with controls, which was consistent with their lower proliferation *in vitro* **(Figure 7A)**. Moreover, controls and glycoengineered cells exhibited similar invasion patterns limited to the superior layer of the intervening mesenchyme (mesoderm) and an overall diffuse morphology **(Figure 7A)**, which was also observed in wild type control cells (data not shown). Interestingly, STn overexpression did not impact proliferation but significantly increased invasion *in vitro* **(Figure 7B)**, in clear contrast with *C1GALT1* KO. These cells also presented increased capacity to form colonies in semisolid substrates and enhanced chemoresistance. However, according to CAM assays, these cells formed smaller tumours without displaying significant alterations in invasion compared to controls. Nevertheless, the tumours exhibited a less cohesive phenotype characterized by high number of isolated tumour niches, which may be related to its higher invasion capacity demonstrated *in vitro*. Finally, *GCNT1* KO showed no evident changes in proliferation, invasion, capacity to form colonies, and chemoresistance *in vitro* **(Figure 7C)**. However, in CAMs, *GCNGT1* KO models invaded more, reaching the lower allantoic epithelium (endoderm) and, in some cases, even expanding beyond that. In summary, we have highlighted the different contributions of distinct glycosylation patterns associated with hypoxia and glucose suppression to cancer progression and potentially dissemination. Most immature glycoforms had major impact on cell proliferation, as previously suggested by us and other groups^81,87^. These cells also showed higher metastatic potential as well as higher tolerance to cisplatin, in agreement with observations made for similar models from other solid tumours^88–90^. Notably, increased sialylation and consequent STn expression promoted cell detachment and formation of less cohesive tumours, reinforcing the close link between this antigen and enhanced cell motility^12,80,81,91^. Finally, cells predominantly expressing sialylated and fucosylated T antigen are markedly more invasive *in vivo*, without other major functional alterations. Building on available literature, we hypothesize that changing the glycosylation of membrane receptors may decisively impact on relevant downstream oncogenic pathways^1,92,93^. We are currently trying to identify common molecular grounds between hypoxic and glucose deprived cells and simple cell models that may account for these observations, which will be decisive to support future targeted therapeutic interventions.

**7.**
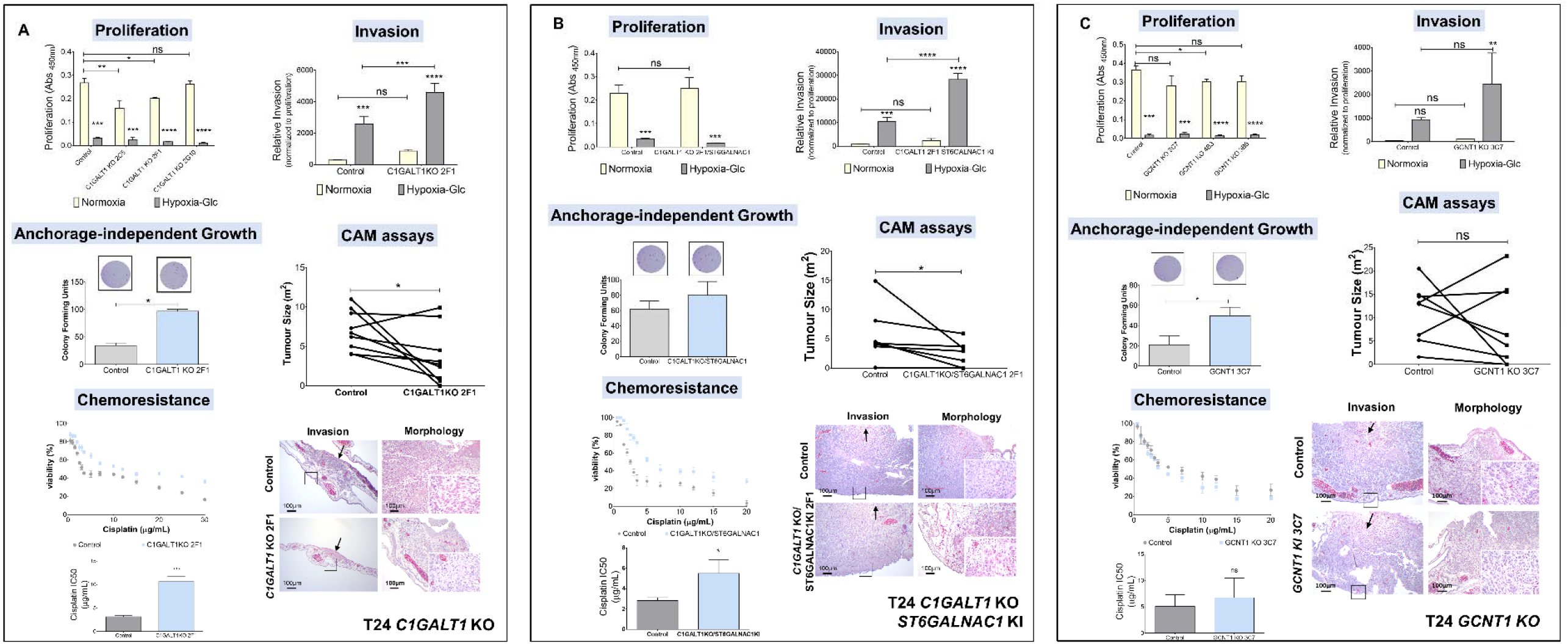
Hypoxia and low-glucose associated glycophenotypes have major functional implications with impact in relevant cancer-associated hallmarks. Cells displaying a marked immature glycophenotype composed by Tn and STn antigens and without further *O*-glycans extension (T24 *C1GALT1* KO **(A)** and *C1GALT1* KO/*ST6GALNAC1* KI **(B)**) showed decreased proliferation *in vitro* and in *in vivo* CAM tumours, higher capacity to form colonies in soft agar, suggesting increased anoikis resistance, and higher tolerance to cisplatin. Both models also showed decreased proliferation and higher invasion *in vitro* under hypoxia and low glucose. These cell lines also revealed a trend increase in invasion in relation to controls *in vitro*, which was not evident in CAM tumours. Moreover, T24 *C1GALT1* KO/*ST6GALNAC1* KI cells formed less cohesive tumours, suggesting increased cell motility. The T24 *GCNT1* KO models overexpressing ST and STn did not show relevant alterations in proliferation *in vitro* or in *in vivo* under normoxia, did not increase the number of CFUs nor chemoresistance. However, it showed increased invasion capacity *in vitro* under hypoxia as well as *in vivo*. The arrows in CAM panels point inoculation areas and squares highlight invasion fronts. ns: not significative; \**p* < 0.05; \*\**p* < 0.01; \*\*\**p* < 0.001; **** *p* < 0.0001 (two-way ANOVA Tukey post hoc test for proliferation and invasion assays *in vitro*; Mann-Whitney for CFU assays; Wilcoxon signed-rank test for CAM tumour size).

## 4. Concluding remarks

Hypoxia and glucose deprivation are salient features of solid tumours arising from sustained proliferative signalling and defective neoangiogenesis. Frequently, cells in the tumour core are faced with limited oxygen and glucose supplies, which decisively shapes their molecular and functional phenotypes. However, low glucose settings have been mostly linked to uncontrolled cell proliferation in the presence of oxygen (the Warburg effect). Furthermore, the contribution of hypoxia to disease severity, even though extensively studied and rather well known, has been addressed without considering the influence of low glucose. Acknowledging the complex microenvironmental challenges experienced by cells striving to survive in low vascularized areas, the present study devoted to understanding the molecular and functional adaptations experienced by cancer cells under low oxygen and glucose to bladder cancer. Namely, we have attempted to comprehensively portrait the impact of these microenvironmental stressors at the transcriptome and metabolome levels, which ultimately led us to comprehensively interrogate the glycome and its biological implications in disease. Our goal was to better understand bladder cancer cells plasticity and adaptability to selective pressures, providing means for more educated and precise interventions.

Amongst the most striking observations arising from this study was that bladder cancer cell lines of distinct genetic and molecular backgrounds responded similarly to microenvironmental cues, denoting common biological grounds facing stress. Bladder cancer cells tolerated well environmental stress, maintaining viability, and avoiding apoptosis, while dramatically decreasing proliferation. This was accompanied by higher invasive capacity, denoting an active strategy to escape to suboptimal microenvironments. Cells under stress also showed increased capacity to tolerate cisplatin, commonly used in the clinics against non-proliferative bladder cancer cells, possibly explaining the resilient nature of poorly vascularized bladder tumours^69,94^. Interestingly, this aggressive behaviour was completely reverted by re-oxygenation and glucose restoration, showing the adaptive capacity of bladder cancer cells and the rapid accommodation to microenvironmental cues. These findings provide a decisive link between the events underlying poor tumour vascularization and the promotion of cancer aggressiveness and were directly linked to major transcriptome and metabolome remodelling. We concluded that, under stress, bladder cancer cells adopted *β*-oxidation rather than anaerobic glycolysis as main bioenergetic pathway, explaining the tremendous decrease in lactate production by these cells. Similar observations were made for prostate cancer, with the fatty acid metabolism rather than glycolysis being suggested as potential basis for imaging diagnosis and targeted treatment^95^. To some extent, this also challenges the pivotal role played by lactate in the aggressiveness of hypoxic cancer cells suggested by many studies^96,97^. We have also found strong indications of mitochondrial autophagy, which has been previously described as a concomitant cancer survival mechanism under nutrient deprivation^98,99^. These adaptions allowed cancer cells to support low energy requirement and cellular basal functions and are consistent with drastic stop in proliferation. Interestingly, while more proliferative bladder tumours are generally more aggressive, they are also more prone to harbour hypoxic niches as result of poor vascularization, which may play a decisive role driving disease fate^100^.

Concomitant to major transcriptome and metabolome reprogramming, we observed major alterations in mucin-type *O*-glycans biosynthesis that may implicate changes in the glycosylation of membrane proteins with functional implications. Glycan biosynthesis was significantly antagonized by glucose limitation, which was further aggravated by the introduction of hypoxia. As a result, bladder cancer cells exhibited more immature glycoforms, namely the Tn antigen and, to less extent, STn and core 3. Some subpopulations also co-expressed increased levels of ST antigen, rather than more extended glycans. Notably, these cells significantly downregulated *C1GALT1C1*, essential for T-synthase (*C1GALT1*) function and glycan chains elongation beyond the core 1 structure. Interestingly, according to the Human Protein Atlas (http://www.proteinatlas.org), low expression of *C1GALT1C1* associates with poor prognosis in bladder cancer (*p*=0.038). However, we found no evidence of decreased *COSMC* or *C1GALT1* to support this as the main mechanism for the expression of immature *O*-glycans by hypoxic and glucose deprived cells. Therefore, we believe that this effect may be intimately linked to nucleotide sugars shortage dictated by low glucose, which could be potentially aggravated by a net disorganization of secretory pathways. Overall, cells under stress acquired simple cell glycophenotypes that are well known to be implicated in an onset of bladder cancer hallmarks, such as invasion, immune escape, and metastasis development^1^. According to observations in patient samples, these glycoforms are more often overexpressed in advanced stage tumours, frequently linked to poor prognosis, but not in the healthy urothelium, highlighting their cancer-related nature and association with disease severity^12,81–83^.

Furthermore, the STn antigen has been found in less proliferative tumour areas showing high HIF-1α content, in accordance with studies *in vitro*^12,81^. Also, experiments involving patient-derived xenografts have suggested a close link between the presence of immature *O*-glycans and the microenvironment^101^, which has finally been demonstrated in the present study. Finally, functional studies implicating glycoengineered cell models have provided decisive evidence that immature glycomes are key drivers of cancer aggressiveness linked to hypoxia and glucose deprivation. We have shown that these molecular alterations are part of a wide array of responses that make bladder cancer cells more capable of tolerating microenvironmental stress; namely, by enabling cells to stop proliferation, resist anoikis, invade *in vivo* and tolerate chemotherapy. Moreover, it provides escape mechanisms, endowing cancer cells with less cohesive and invasive traits. These observations are in perfect agreement with other reports from simple cells cancer models of different origins, highlighting the pancarcinomic nature of these alterations. Furthermore, it provides novel grounds to understand the events underlying alterations in glycosylation, which have so far been scarcely linked to functional mutations in *COSMC*^102^ and, more recently, with pro-inflammatory immune responses^103^.We also noted the tremendous plasticity of *O*-glycosylation pathways facing re-oxygenation and restored access to glucose, as observed at other levels. Collectively, we hypothesize that alterations in glycosylation hold potential to identify cells lying dormant in hypoxic niches, that may eventually be responsible by recapitulating the disease upon reoxygenation.

In summary, this study was able to provide the microenvironmental context for previous observation regarding mucin-type *O*-glycans expression in bladder tumours, with potential implications for other solid tumours where short-chain glycans also play key functional roles^104^. Since most of these glycoepitopes, such as Tn and STn antigens, are not expressed by the healthy urothelium, they also provide a unique opportunity for precise cancer targeting. Namely, it offers a strategy to detect and target hypoxic cells that, as shown by this and other studies^1,104^, constitute more aggressive subpopulations of cancer cells with limited therapeutic options.

## Supporting information

Peixoto et al. 2021 preprint Supp. Information

## Acknowledgments and Funding Sources

The authors wish to acknowledge the Portuguese Foundation for Science and Technology (FCT) for the human resources grants: PhD grant, SFRH/BD/111242/2015 (AP), 2020.08708.BD (RF), 2020.09384.BD (DF), SFRH/BD/146500/2019 (MRS), SFRH/BD/127327/2016 (CG), SFRH/BD/142479/2018 (JS), and FCT assistant researcher grant CEECIND/03186/2017 (JAF). FCT is co-financed by European Social Fund (ESF) under Human Potential Operation Programme (POPH) from National Strategic Reference Framework (NSRF). The authors also acknowledge FCT funding for the CI-IPOP research unit (PEst-OE/SAU/UI0776/201) and the LAQV research unit (UIDB/50006/2020), the Portuguese Oncology Institute of Porto Research Centre (CI-IPOP-29-2016-2020, CI-IPOP-58-2016-2020, CI-IPOP-Proj.70-bolsa2019-GPTE), and the ICBAS PhD Program in Biomedical Sciences. This article is also a result of the project NORTE-01-0145-FEDER-000012, supported by Norte Portugal Regional Operational Programme (NORTE 2020), under the PORTUGAL 2020 Partnership Agreement, through the European Regional Development Fund (ERDF). This work was financed by FEDER - Fundo Europeu de Desenvolvimento Regional funds through the COMPETE 2020 - Operacional Programme for Competitiveness and Internationalisation (POCI), Portugal 2020, and by Portuguese funds through FCT/MCTES - Fundação para a Ciência e a Tecnologia/Ministério da Ciência, Tecnologia e Ensino Superior in the framework of the project "Institute for Research and Innovation in Health Sciences" (POCI-01-0145-FEDER-007274). Finally, the authors wish to acknowledge the support of the i3S scientific platforms for the *In vivo* CAM assays and HEMS, member of the national infrastructure PPBI - Portuguese Platform of Bioimaging (PPBI-POCI-01-0145-FEDER-022122). Moreover, the authors acknowledge the kind collaboration of the pharmaceutical units of IPO-Porto for providing and supervising the handling of clinical grade cisplatin.

## Author Contributions

JAF conceived and supervised the study; AP, MRS, AMNS performed glycomics analysis, AP, DF, CP, RF, and CG performed immunoassays; MC and PP aided AP with genetic editing; AP, MRS, JS, FT, JAF and AMNS curated the data, performed bioinformatics, and statistical analysis; RF and MJO gave valuable scientific input and provided technological resources for project execution; LLS and JAF gathered the financial and human resources for the study; AP and JAF wrote the manuscript, which was reviewed and commented by all the co-authors.

