## Supplementary material for "Metabolomics, Transcriptomics and Functional Glycomics Reveals Bladder Cancer Cells Plasticity and Enhanced Aggressiveness Facing Hypoxia and Glucose Deprivation": Peixoto et al. 2021 preprint Supp. Information

**-Supporting Information-**

Andreia Peixoto^1,2,3,4,5^, Rui Freitas^1,2,3,4^, Dylan Ferreira^1,2^, Marta Relvas-Santos^1,2,3,4,6^, Cristiana Gaiteiro^1,2^, Paula Paulo^7^, Marta Cardoso^7^, Janine Soares^1,2,8^, Carlos Palmeira^1,9,10^, Filipe Teixeira^6^, Rita Ferreira^8^, Maria José Oliveira^3,4^, André M. N. Silva^6^, Lúcio Lara Santos^1,2,5,10,11^ José Alexandre Ferreira^1,2,5^

^1^Experimental Pathology and Therapeutics Group, IPO Porto Research Center (CI-IPOP), Portuguese Oncology Institute, 4200-072 Porto, Portugal; ^2^Institute of Biomedical Sciences Abel Salazar (ICBAS), University of Porto, 4050-313 Porto, Portugal; ^3^Institute for Research and Innovation in Health (i3S), University of Porto, 4200-135 Porto, Portugal; ^4^Institute for Biomedical Engineering (INEB), University of Porto, 4200-135 Porto, Portugal; ^5^Porto Comprehensive Cancer Center (P.ccc), 4200-072 Porto, Portugal; ^6^REQUIMTE-LAQV, Department of Chemistry and Biochemistry, Faculty of Sciences of the University of Porto, 4169-007 Porto, Portugal; ^7^Cancer Genetics Group, IPO Porto Research Center (CI-IPOP), Portuguese Oncology Institute, 4200-072 Porto, Portugal; ^8^REQUIMTE-LAQV, Department of Chemistry, University of Aveiro, 3810-193 Aveiro; ^9^Immunology Department, Portuguese Institute of Oncology of Porto, 4200-072 Porto, Portugal; Portugal;^10^Health School of University Fernando Pessoa, 4249-004 Porto, Portugal; ^11^Department of Surgical Oncology, Portuguese Oncology Institute, 4200-072 Porto, Portugal.

**Corresponding author:**

José Alexandre Ferreira

Experimental Pathology and Therapeutics Group, Research Centre, Portuguese Oncology Institute of Porto, R. Dr. António Bernardino de Almeida 62, 4200-072 Porto, Portugal; Tel. +351 225084000 (ext. 5111).

**Running head:** Bladder Cancer Cells Plasticity in Hypoxia and Low Glucose

**Keywords:** glycomics; metabolomics; transcriptomics; bladder cancer; microenvironment; hypoxia

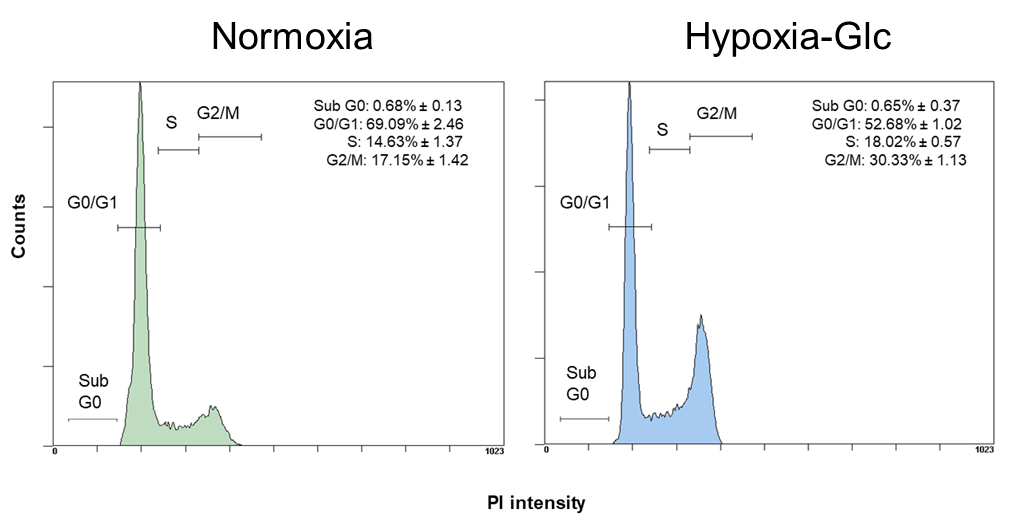

**Figure S1. Hypoxia and low glucose induce an arrest of cell cycle in G2/M phases.** The histograms highlight the distribution of bladder cancer cells in normoxia and hypoxia and low glucose according to their cell cycle. This clearly shows an increase in the number of cells in G2/M under microenvironmental stress.

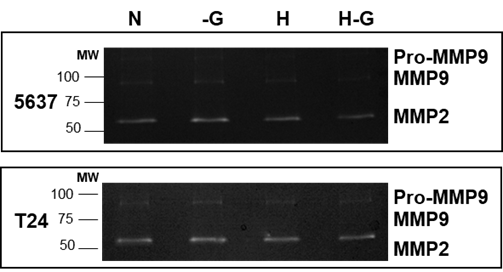

**Figure S2. Bladder cancer cells under hypoxia and low glucose do not display major changes in proteolytic activity.** The panel highlights the no changes in proteolytic activity, herein translated by MMP2 and 9 that are two major metalloproteinases in bladder cancer, in low glucose (-Glc), hypoxia and the combination of both factors.

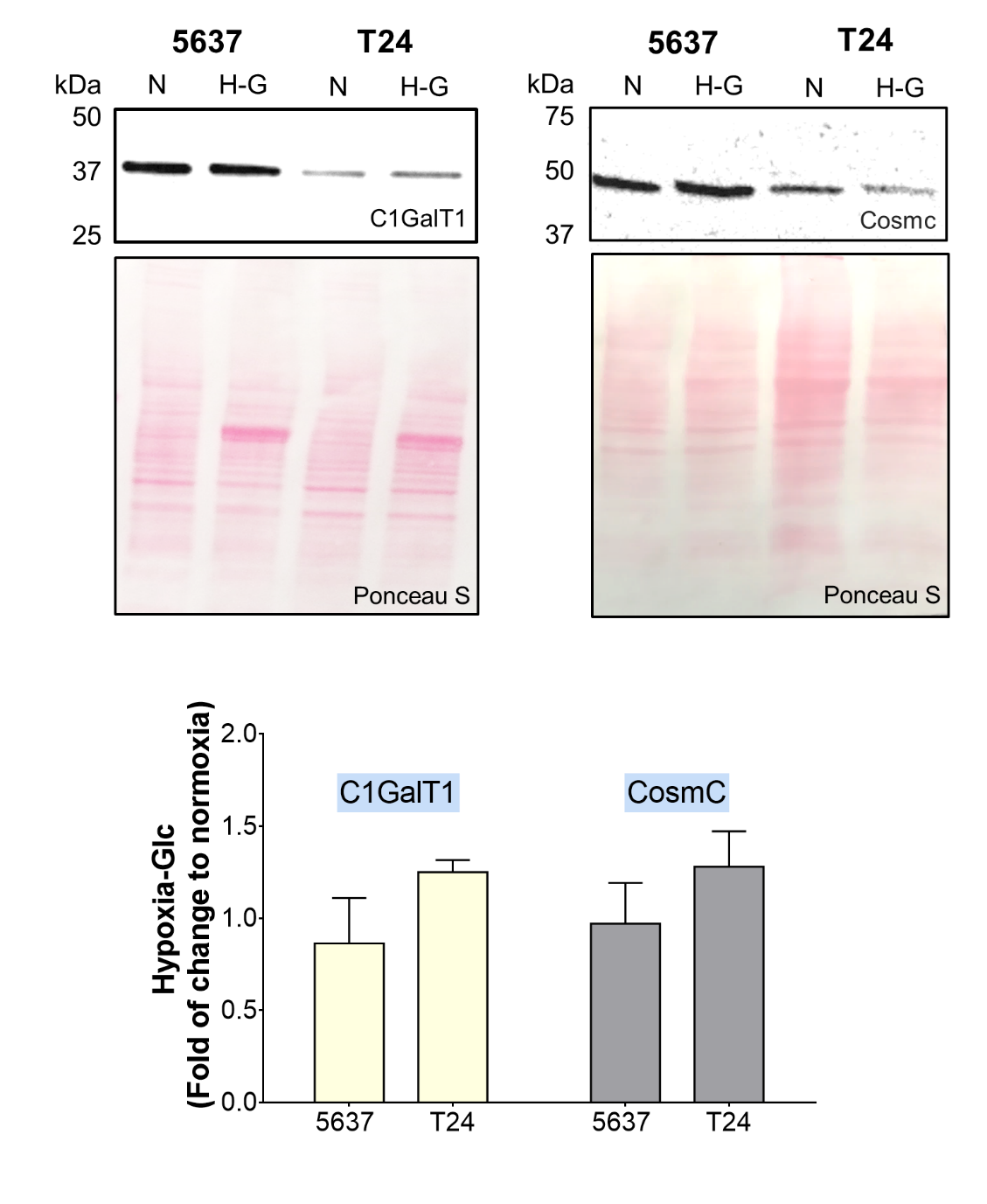

**Figure S3. C1GalT1 and its chaperone Cosmc expressions do not change significantly with hypoxia and low glucose.**

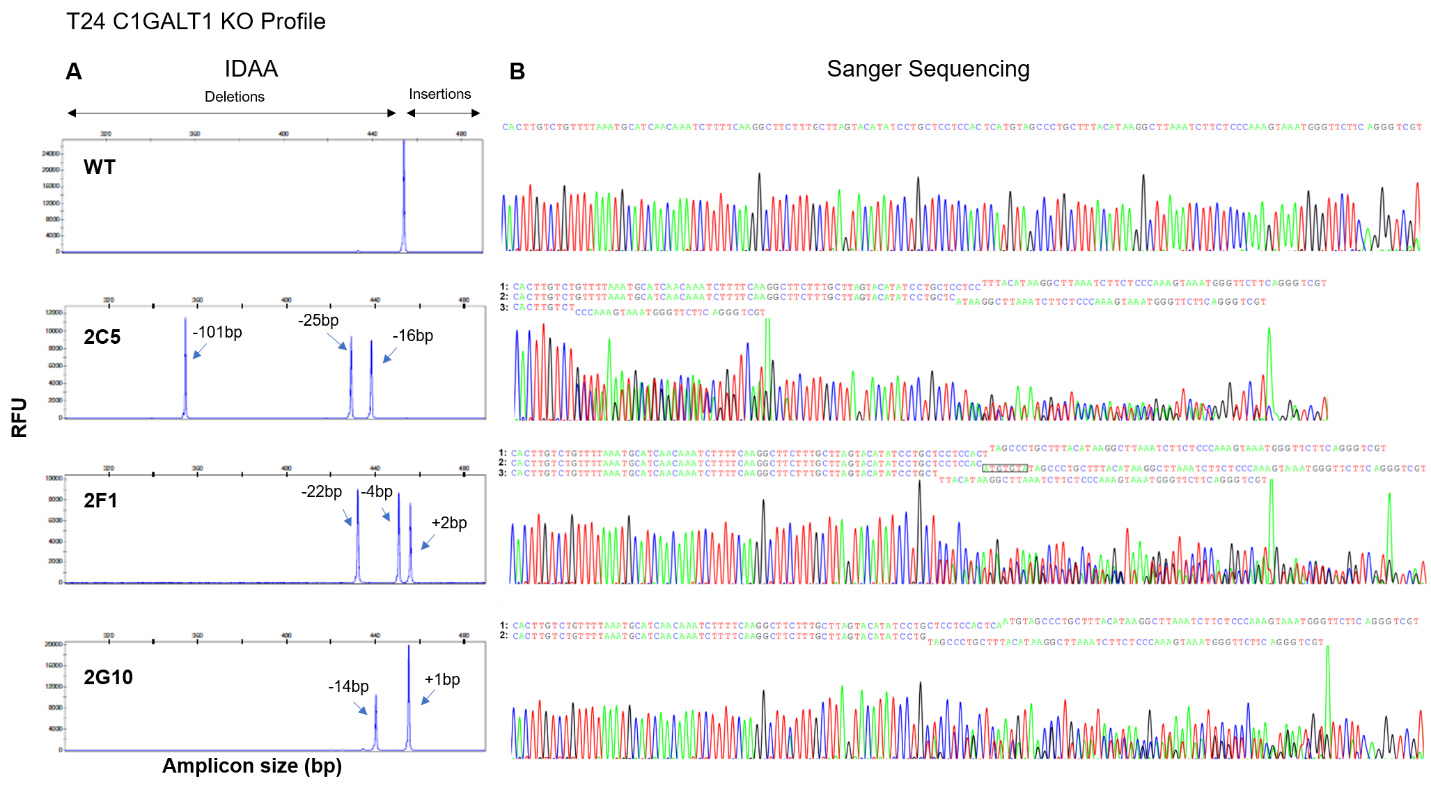

**Figure S4.** **Genomic profiling of glycoengineered T24 *C1GALT1* KO models.** T24 cells were glycoengineered to knock-out human C1GALT1 and three clones with diverse indels were selected for proof-of-concept experiments. Induced indel mutations were characterized by Indel Detection by Amplicon Analysis (IDAA) **(A)** and Sanger Sequencing **(B)**. Electropherograms of Sanger Sequencing with the reverse primer are shown, along with the corresponding sequencing readings (1, 2 and 3). Accordingly, the 2C5 clone is characterized by three different DNA sequences, with the following observed variants and predicted consequences: **1:** a c.610_625del [p.(Gln204GlufsTer7)], leading to a 16bp deletion and the replacement of Gln 204 by Glu, resulting in a frame-shift introducing a stop codon 7 a.a. ahead; **2:** a c.605_629del [p.(Val202GlufsTer6)], resulting in a 25bp deletion and the replacement of Val 202 by Glu, leading to a frame-shift introducing a stop codon 6 a.a. ahead; and **3:** a c.589_689del [p.(Arg197GlnfsTer5)], in which a 101bp deletion results in the replacement of Arg 197 by Gln, leading to a frame-shift introducing a stop codon 5 a.a. ahead. Similarly, clone 2F1 is also characterized by three different DNA sequences, with the following observed variants and predicted consequences: **1:** a c.618_621del [p.(Tyr206Ter)], where a 4bp deletion results in the replacement of Tyr 206 by a stop codon; **2:** a c.618_622delinsTACACAT [p.(Met207ThrfsTer10)], where a 5bp deletion and a 7bp insertion results in the replacement of Met 207 by Thr, inducing a frame-shift and introducing a stop codon 10 a.a. ahead; and **3:** a c.614_635del [p.(Gly205AspfsTer4)], where a 22bp deletion results in the replacement of Gly 205 by Asp, leading to a frame-shift introducing a stop codon 4 a.a. ahead. Finally, clone 2G10 is characterized by two different DNA sequences, with the following observed variants and predicted consequences: **1:** a c.620dup [p.(Met207IlefsTer19)], where the duplication of nucleotide 620 results in the replacement of Met 207 by Ile, leading to a frame-shift introducing a stop codon 19 a.a. ahead; and **2:** a c.620_633del [p.(Met207ArgfsTer14)], resulting in a 14bp deletion and the replacement of Met 207 by Arg, resulting in a frame-shift introducing a stop codon 14 a.a ahead.

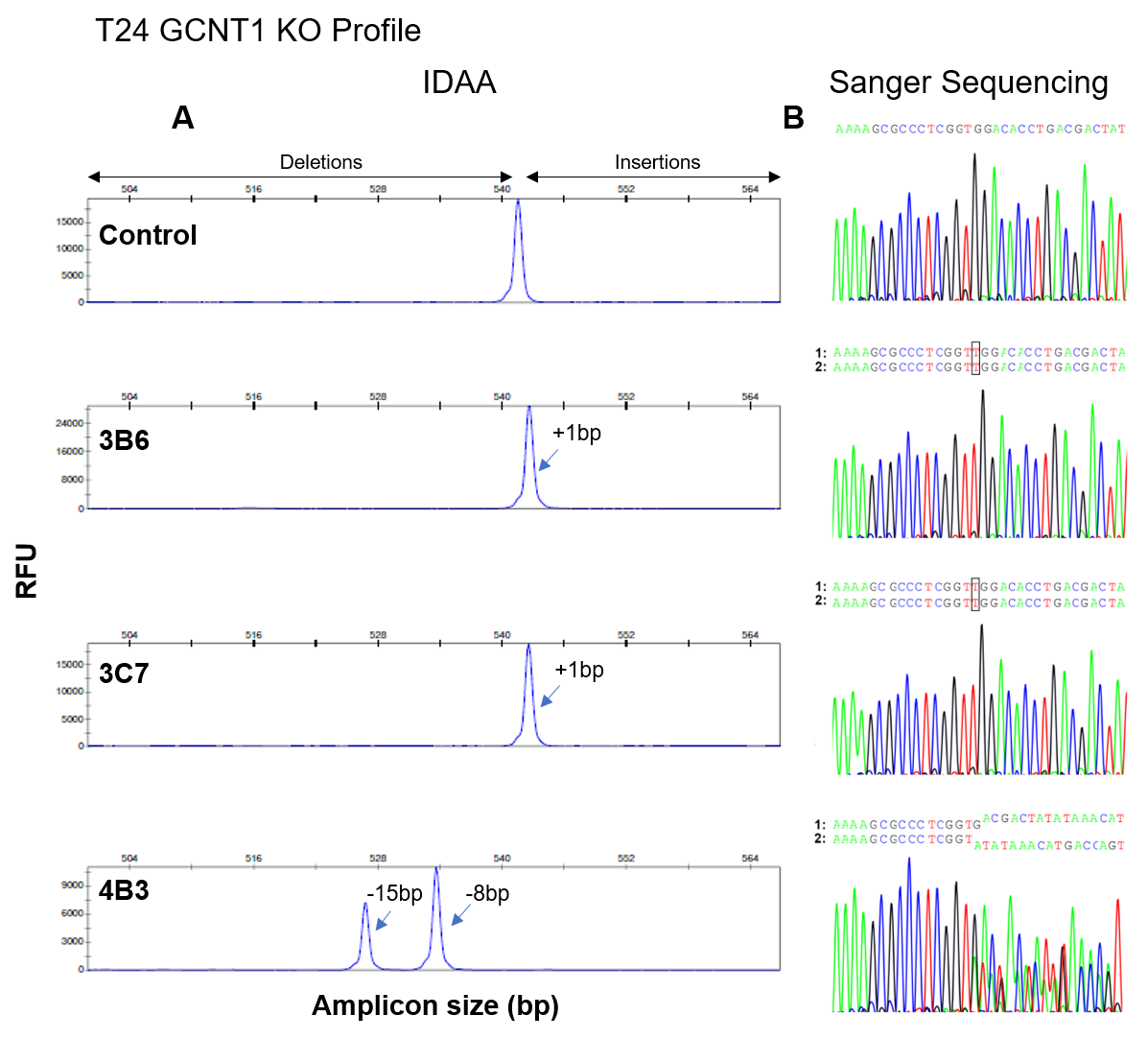

**Figure S5.** **Genomic profiling of glycoengineered T24 *GCNT1* KO models.** T24 cells were glycoengineered to knock-out human GCNT1 and three clones were selected for proof-of-concept experiments. Induced indel mutations were characterized by Indel Detection by Amplicon Analysis (IDAA) **(A)** and Sanger Sequencing **(B)**. Electropherograms of Sanger Sequencing with the forward primer are shown, along with the corresponding sequencing readings (1 and 2). Accordingly, both 3B6 and 3C7 where characterized by a single DNA sequence reading (assumed homozygous), with the c.262dup [p.(Trp88LeufsTer4)] being observed, where duplication of nucleotide 262 is predicted to lead to the replacement of Trp 88 by Leu, leading to a frame-shift introducing a stop codon 4 a.a ahead. In turn, 4B3 clone is characterized by two different DNA sequences, with the following observed variants and predicted consequences: **1:** a c.264_271del [p.(Trp88Ter)], where a 8bp deletion results in the replacement of Trp 88 by a stop codon; and **2:** a c.263_277del [p.(Trp88_Asp92del)], where a 15bp in-frame deletion between Trp 88 and Asp 92 results in the suppression of 5 a.a and the generation of a truncated protein.

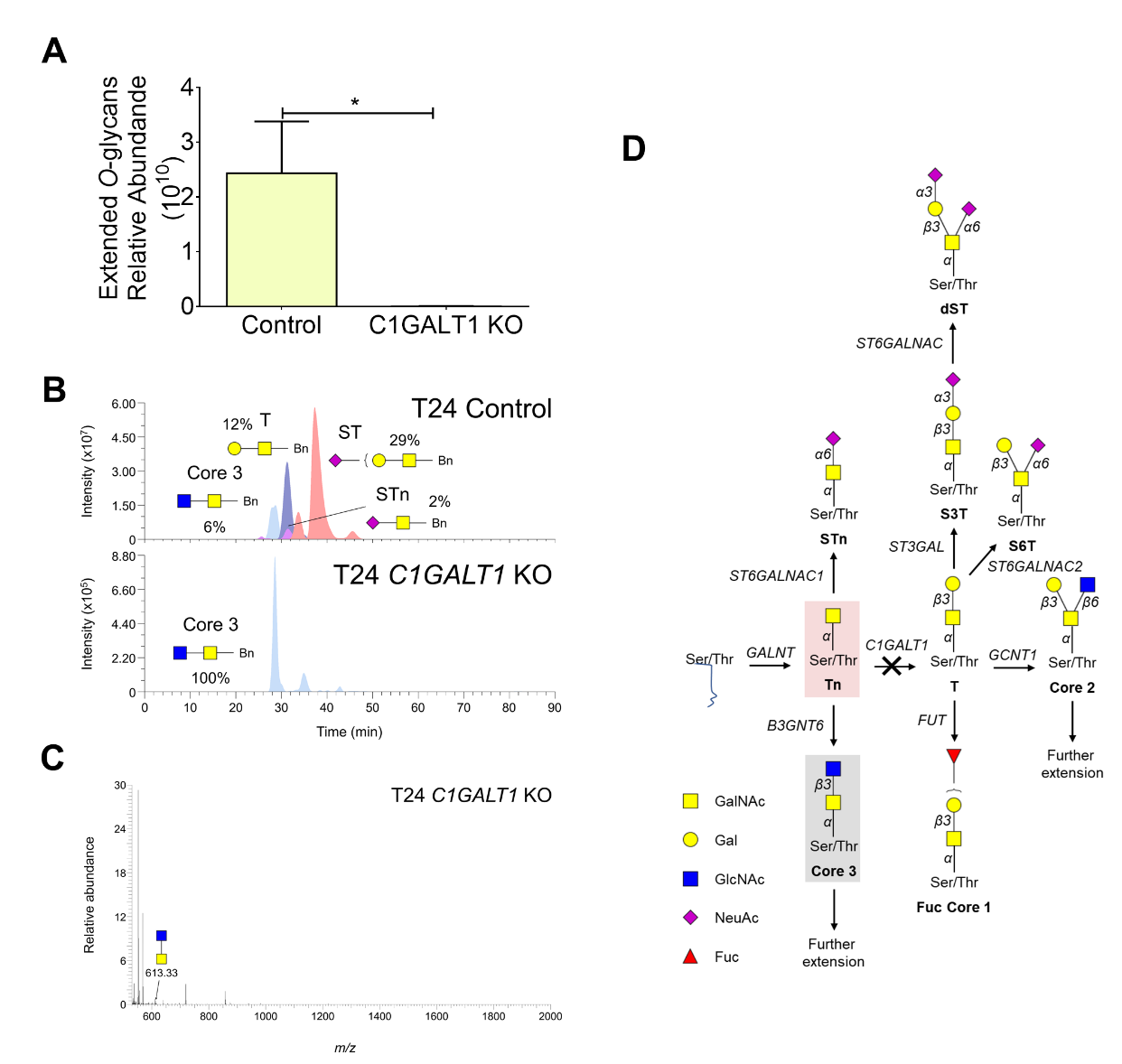

**Figure S6. Mass spectrometry highlights the abrogation of glycans extension and accumulation of immature *O*-glycans after *C1GALT1* KO. A) *C1GALT1* KO promotes a major suppression of extended *O*-glycans in T24 cells.** *C1GALT1* KO significantly reduced the expression of extended *O*-glycans to residual levels in comparison to controls. **B) Extracted ion chromatograms (EICs) and C) full MS spectrum highlighting the loss of extended *O*-glycans such as the T and ST antigens in glycoengeneired T24 cells.** Glycans percentage is expressed in relation to the total glycans identified for these cell lines by MS/MS. Panel B shows ECIs for *O*-glycans characteristic of hypoxic and low glucose grown cells (*i.e.* core 3, STn, T, and ST). The *O*-glycome of control cells presented high percentage of ST and, to less extent, T antigens. Low amounts of core 3 and STn were also present. *C1GALT1* KOs only presented core 3 as result of significant decrease in extended *O*-glycans. **D) Schematic representation of *O*-glycans pathways highlighting the glycans expressed by *C1GALT1* KOs, according to MS/MS analysis and immunoassays (Figure 6-main manuscript).** MS analysis shows the loss of extended *O*-glycans, whereas immunoassays confirm the expression of also low amounts of core 3 and, potentially, STn. Light pink: Main expressed glycan; Light grey: less abundant, immature glycans.

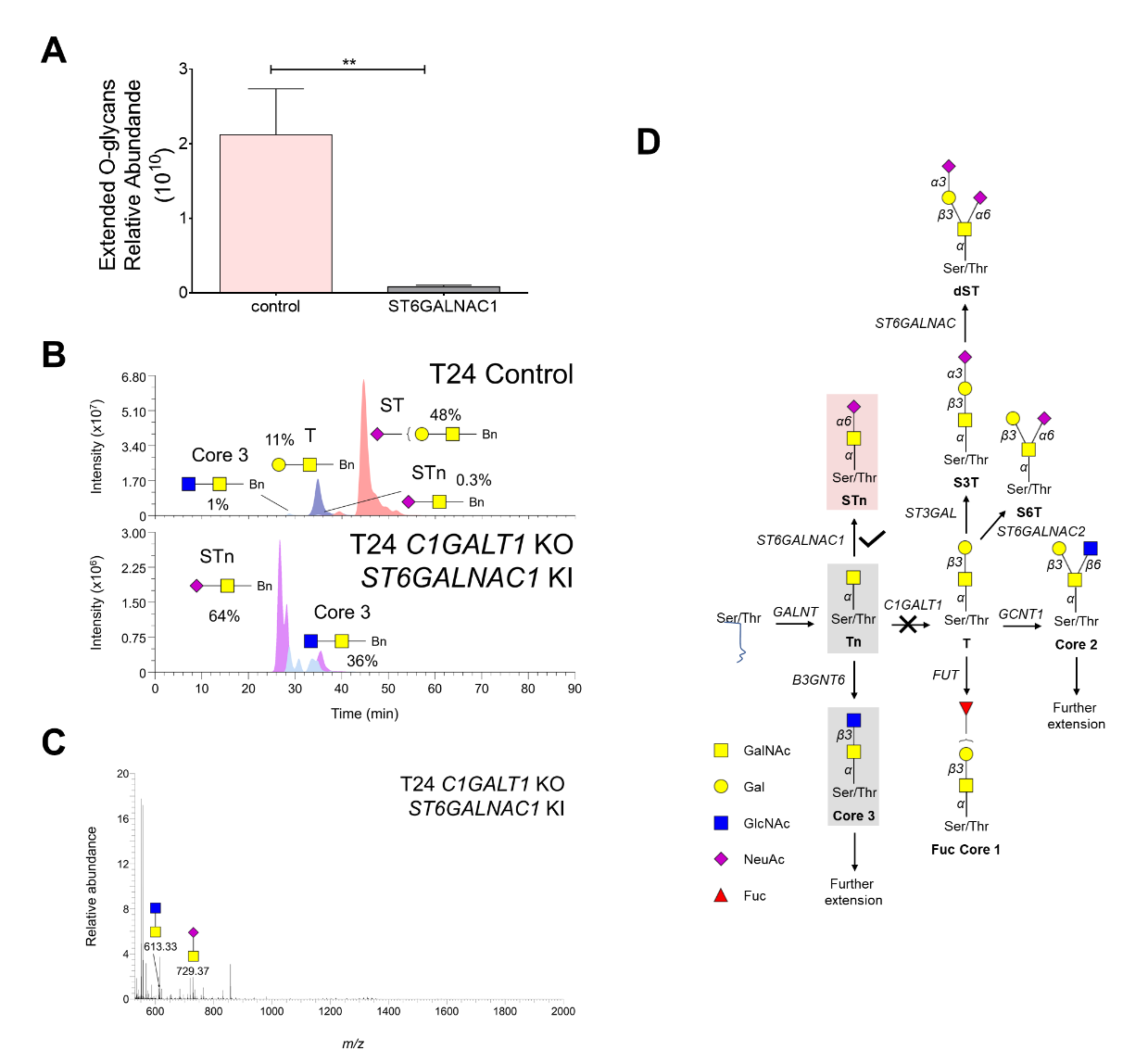

**Figure S7. Mass spectrometry demonstrates overexpression of STn after *ST6GALNAC1* KI in T24 *C1GALT1* KO cells. A) *ST6GALNAC1* overexpression promotes a major increase in STn in T24 C1GALT1 cells.** STn overexpression significantly reduced the percentage of extended *O*-glycans to residual levels in comparison to controls and parental cells. **B) Extracted ion chromatograms (EICs) and C) full MS spectrum highlighting the loss of extended *O*-glycans such as the T and ST antigens and the overexpression of STn in glycoengineered T24 cells.** Glycans percentage is expressed in relation to the total glycans identified for these cell lines by MS/MS. Panel B shows ECIs for *O*-glycans characteristic of hypoxic and low glucose grown cells (*i.e.* core 3, STn, T, and ST). The *O*-glycome of control cells presented high percentage of ST and, to less extent, T antigens. Low amounts of core 3 and STn were also present. *C1GALT1* KO/ST6GALNAC1 KI cells mostly presented STn, low amounts of Tn and core 3, explaining the tremendous decrease in extended *O*-glycans. **D) Schematic representation of *O*-glycans pathways highlighting the glycans expressed by *C1GALT1* KO/ST6GALNAC1 KI cells, according to MS/MS analysis and immunoassays (Figure 6-main manuscript).** MS analysis shows the loss of extended *O*-glycans, whereas immunoassays confirm the expression of STn antigens and, to less extent core 3 and Tn. Light pink: Main expressed glycan; Light grey: less abundant, immature glycans.

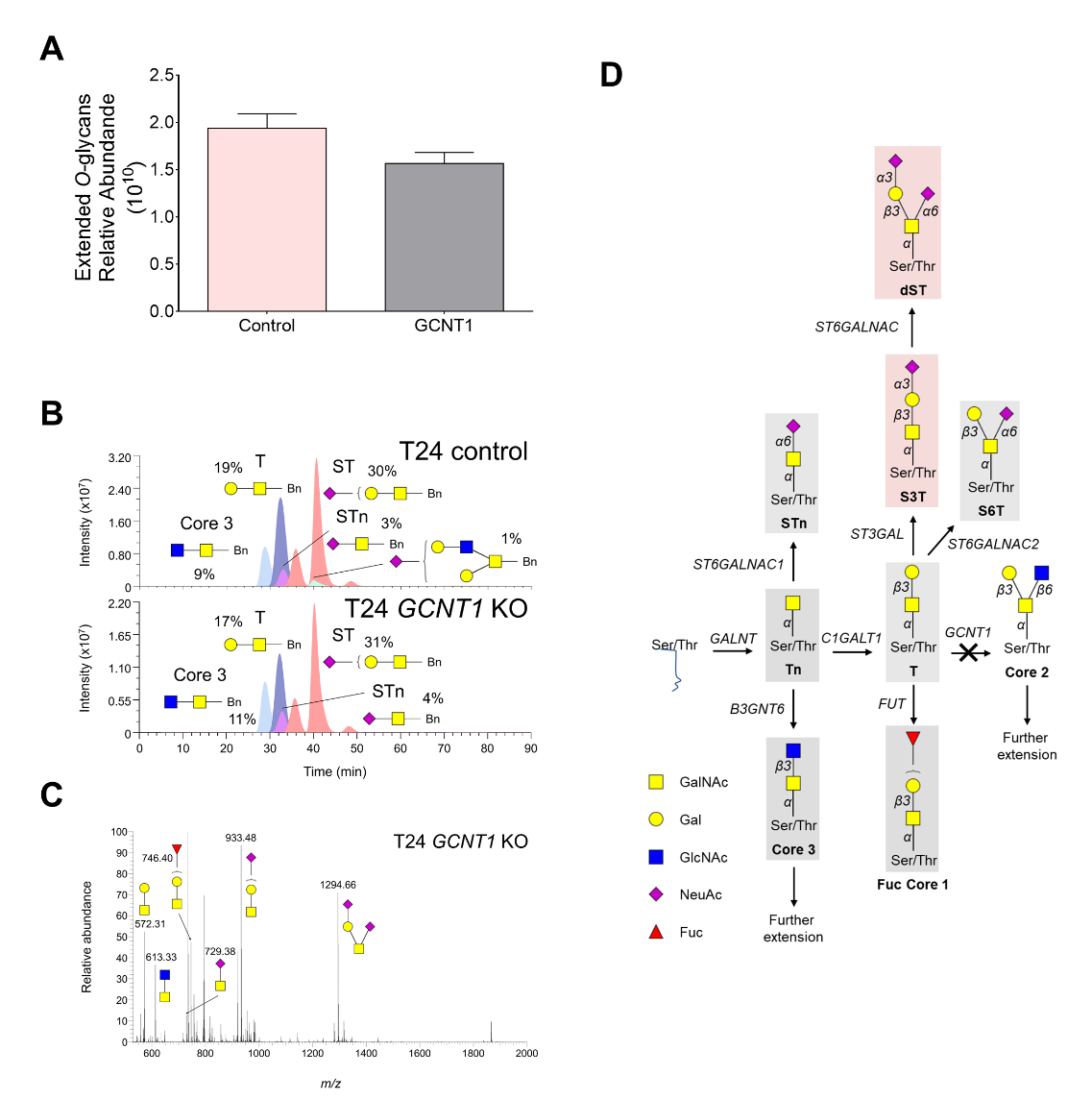

**Figure S8. Mass spectrometry demonstrates that *GCNT1* KO in T24 cells leads to the loss of glycans extended beyond core 1. A) *GCNT1* overexpression promotes a major increase in STn in T24 C1GALT1 cells.** *GCNT1* KO led to a trend decrease in the expression of extended *O*-glycans in comparison to controls, consistent with the loss of core 2 derived glycans. **B) Extracted ion chromatograms (EICs) and C) full MS spectrum highlight that *GCNT1* KO glycome is mainly composed by sialylated core 1 structures and, to less extent, fucosylated T antignes.** Glycans percentage is expressed in relation to the total glycans identified for these cell lines by MS/MS. Panel B shows ECIs for *O*-glycans characteristic of hypoxic and low glucose grown cells (*i.e.* core 3, STn, T, and ST). The *O*-glycome of control cells presented high percentage of ST and, to less extent, T antigens. Low amounts of core 3 and STn were also present. *GCNT1* KI cells mostly presented sialylated T antigens and fucosylated core 1 as major ions. The STn and core 3 antigens were also detectable. No evidences of *O*-glycans extended beyond core 1 were observed. **C) Schematic representation of *O*-glycans pathways highlighting the glycans expressed by *GCNT1* cells, according to MS/MS analysis and immunoassays (Figure 6-main manuscript).** MS analysis shows that T24 do not extend *O*-glycans beyond core 1 and the predominance of sialylated T antigens, whereas immunoassays confirm the expression of ST antigens and, to less extent STn and core 3. Light pink: Main expressed glycan; Light grey: less abundant, immature glycans.
